# Task-irrelevant stimuli reliably boost phasic pupil-linked arousal but do not affect decision formation

**DOI:** 10.1101/2024.05.14.594080

**Authors:** J. Hebisch, A.-C. Ghassemieh, E. Zhecheva, M. Brouwer, S. van Gaal, L. Schwabe, T.H. Donner, J.W. de Gee

## Abstract

The arousal systems of the brainstem, specifically the locus coeruleus-noradrenaline system, respond “phasically” during decisions. These central arousal transients are accompanied by dilations of the pupil. Mechanistic attempts to understand the impact of phasic arousal on cognition would benefit from the ability to experimentally manipulate arousal in a temporally precise manner. Here, we evaluated a non-invasive candidate approach for such a manipulation in humans: presenting task-irrelevant auditory stimuli at different latencies during the execution of a challenging task. Task-irrelevant auditory stimuli drive responses of brainstem nuclei involved in the control of pupil size. But it is unknown whether such sound-evoked responses mimic the central arousal transients evoked during cognitive computations. A large body of evidence has implicated central arousal transients in a bias reduction during challenging perceptual decisions. We thus used challenging visual decisions as a testbed, combining them with task-irrelevant sounds of varying onset latency or duration. Across three experiments, the sounds consistently elicited well-controlled pupil responses that superimposed onto task-evoked responses. While we replicated a negative correlation between task-evoked pupil responses and bias established in previous work, the task-irrelevant sounds had no behavioral effect. This dissociation suggests that cognitive task engagement and task-irrelevant sounds may recruit distinct neural systems contributing to the control of pupil size.

## Introduction

Brainstem arousal systems, including the locus coeruleus noradrenaline system, are transiently (“phasically”) active during the performance of challenging cognitive tasks (Aston-Jones & Cohen, 2005; Bouret & Sara, 2005; Breton-Provencher et al., 2022; Breton-Provencher & Sur, 2019; de Gee et al., 2017). Such central arousal transients are likely to play important roles in healthy cognitive behavior (Aston-Jones & Cohen, 2005; Bouret & Sara, 2005; Dayan & Yu, 2006) as well as its aberrations (Arnsten, 2015; Aston-Jones & Cohen, 2005). Therefore, the ability to manipulate central arousal transients in a well-controlled and temporally precise manner could open new opportunities for advancing basic as well as clinical neuroscience.

Dilations of the pupil are a readily assessable peripheral readout of phasic arousal responses and, potentially, experimental manipulations thereof. Non-luminance mediated variations of pupil size track the activity of the locus coeruleus, along with the activity of other subcortical nuclei involved in arousal, orienting, and alerting, such as the cholinergic basal forebrain, dopaminergic midbrain, and the superior and inferior colliculi (Breton-Provencher & Sur, 2019; de Gee et al., 2017; Joshi et al., 2016; Lloyd et al., 2023; Murphy et al., 2014; Varazzani et al., 2015; Wang & Munoz, 2015). Furthermore, the pupil transiently responds (dilates) time-locked to perceptual decisions (Beatty, 1982; Cheadle et al., 2014; Gilzenrat et al., 2010), with a time course that reflects the time course of decision formation (de Gee et al., 2017; de Gee et al., 2014).

Here, we evaluated a promising approach to non-invasively manipulate phasic arousal: task-irrelevant auditory stimuli presented during cognitive tasks. Task-irrelevant sounds can accelerate human responses to (task-relevant) visual stimuli (Hackley et al., 2009; Hackley & Valle-Inclán, 1998; Hershenson, 1962; Jepma et al., 2009; Stahl & Rammsayer, 2005; Tona et al., 2016). Importantly, task-irrelevant sounds dilate the pupil (Cronin et al., 2023; Petersen et al., 2017; Tona et al., 2016) and drive locus coeruleus activity (Grant et al., 1988; Joshi & Gold, 2020, 2022; Joshi et al., 2016). However, other brainstem nuclei also respond to auditory sounds – most strongly the inferior colliculus that is part of the auditory pathway, and whose activity is also read out by pupil size (Joshi et al., 2016). So, it is currently unknown if task-irrelevant sounds can be used to reliably manipulate the same central arousal transients that naturally occur during cognitive behavior.

We focused on challenging perceptual decisions, because a substantial body of evidence implicates phasic arousal in a well-understood behavioral consequence in such decisions: reduction of a choice bias: (i) trial-to-trial variations in the amplitude of task-evoked brainstem and pupil responses due to intrinsic factors (i.e., unrelated to external stimuli or actions) predict a bias reduction across different species and choice tasks (de Gee et al., 2017; de Gee et al., 2020; Krishnamurthy et al., 2017; Lewandowska et al., 2019; Schriver et al., 2020); (ii) systematic variations of the pupil response (due to computational variables such as uncertainty and surprise) affect perceptual decisions in a similar fashion (Krishnamurthy et al., 2017; Murphy et al., 2021; Urai et al., 2017; van den Brink et al., 2023); (iii) a causal link between locus coeruleus activity and choice bias has been established in mice (Breton-Provencher et al., 2022); and (iv) these empirical findings stand on solid grounds of computational theory (Dayan & Yu, 2006; Jordan, 2023; Sales et al., 2019; Yu & Dayan, 2005).

We reasoned that if the central responses to task-irrelevant sounds mimic the central arousal responses occurring naturally during perceptual decisions, then the profile of associated pupil responses, and behavioral correlates should be similar to that of intrinsic task-related phasic arousal. We present three experiments to test this idea.

## Results

We first conducted two parallel experiments in which participants made challenging perceptual decisions based on visual information while we varied the frequency, timing, and duration of an auditory white noise stimulus, henceforth referred to as “task-irrelevant sound” (**Fig. 1**; see **Fig. S1** for task behavior). In Experiment 1, participants reported the presence or absence of a visual Gabor grating of near-threshold contrast superimposed onto flickering visual noise (de Gee et al., 2017). The task-irrelevant sound started together with a baseline interval of 1 s duration (visual noise only) preceding a so-called decision interval, which either contained the Gabor stimulus or not, and the beginning of which was signaled to the participant. The white noise lasted 4 s and occurred only on 25% of trials. Experiment 2 used the same detection task but was tailored to delineate the time course of the effects of the task-irrelevant sound on behavior at high temporal resolution. Here, task-irrelevant sounds were of short duration (100 ms), occurred on 80% of trials, and started at random times between −3 s and 0.5 s with respect to decision interval onset (uniformly distributed).

**Figure 1.**
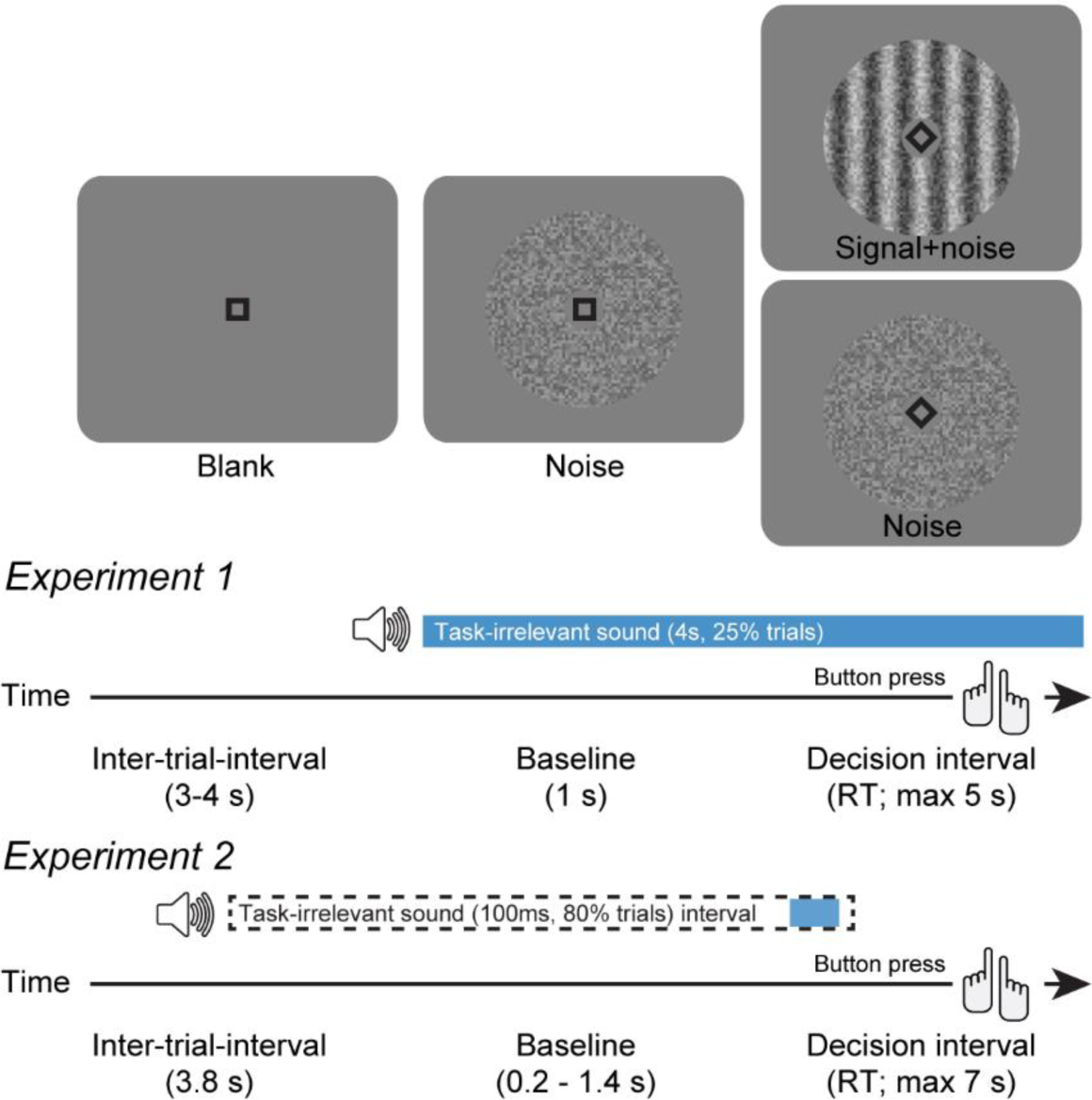
Behavioral tasks. Schematic sequence of events during the simple forced choice task of Experiments 1 and 2. Participants reported the presence or absence of a faint grating signal superimposed onto dynamic noise. Signal contrast is high for illustration only. In Experiment 1, the task-irrelevant, auditory white-noise (70 dB) stimulus started simultaneously to baseline onset (25% of trials). In Experiment 2, brief white noise stimuli (100 ms, 70dB) were played randomly between 3 s before and 0.5 s after decision interval onset (80% of trials).

### Trial-to-trial variability of task-evoked pupil responses

In our experiments, pupil responses during decisions were comprised of a mix of contrast-related constriction at baseline (visual noise) onset (Tsujimura et al., 2001) and a dilatory component related to task engagement that built up during decision formation (Cheadle et al., 2014; de Gee et al., 2017; de Gee et al., 2014). Importantly, we observed substantial trial-to-trial variability in the magnitude of the task-evoked pupil responses (**Fig. 2A**, Methods): bins containing the highest 12.5% of task-evoked pupil responses on trials with no task-irrelevant sound showed a clear dilation (average bin pupil response ± S.E.M: Experiment 1, 18.625 ± 0.154 % signal change; Experiment 2, 19.760 ± 0.292 % signal change) whereas those containing the lowest responses showed pupil constrictions (average bin pupil response ± S.E.M: Experiment 1, −14.342 ± 0.151 % signal change; Experiment 2, −18.548 ± 0.274 % signal change). Because the effect of the presence or absence of the (near-threshold contrast) target signals (superimposed on dynamic noise that was present in all trials, **Fig. 1**, top) on pupil responses is negligible (data not shown), and the responses are locked to response, we conclude that the variability shown in Figure 2A stems from endogenous sources.

**Figure 2.**
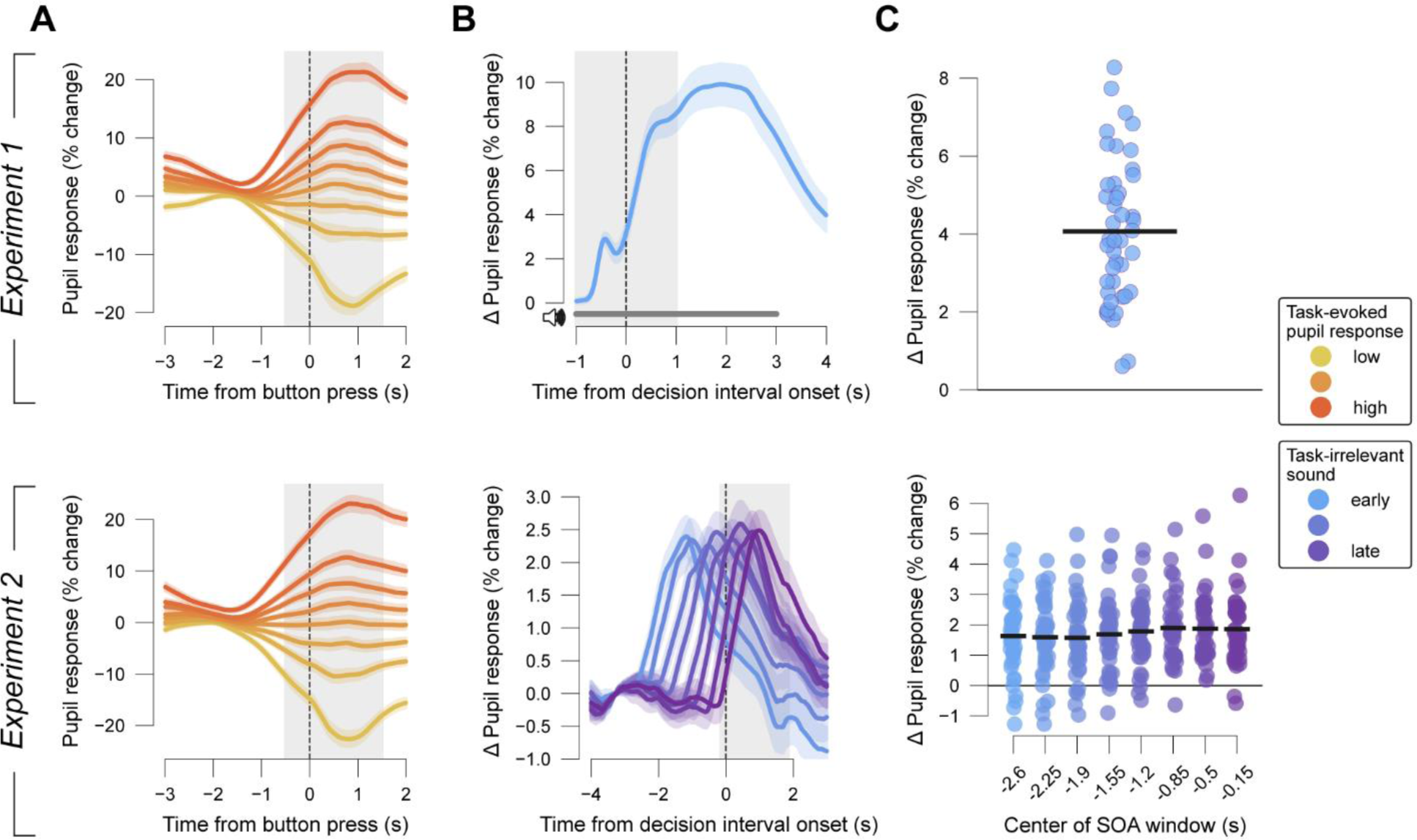
Time courses of pupil responses. **(A)** Pupil response time course on trials with no task-irrelevant sound, binned post-hoc by size of task-evoked pupil response, time-locked to button-press (choice). Shading, S.E.M across participants. Grey shading, interval used for quantifying task-evoked pupil responses (Methods). **(B)** Differential pupil size time courses between task-irrelevant sound trials and trials without task-irrelevant sound. Baselined with 0.5 second intervals from before the first possible task-irrelevant sound. For Experiment 2, data were binned with a window sliding along possible task-irrelevant sound onset asynchronies (SOAs; window size, 800ms; step size 350 ms). Shading, S.E.M across participants. Grey shading, interval used for quantifying task-irrelevant sound-evoked pupil responses (bottom, exemplary for last SOA window centered at −0.15 s; Methods). **(C)** Participant-wise task-irrelevant sound-evoked pupil response magnitude. Black bars, mean across participants.

### Task-irrelevant sounds evoke dissociable and reliable pupil responses

We next quantified the pupil responses specifically evoked by the task-irrelevant auditory white noise stimuli. To this end, we subtracted the average pupil size time course on trials without a task-irrelevant sound from each trial’s time course with a task-irrelevant sound (Methods). The mean resulting time courses showed robust increases in pupil size with respect to pre-trial baseline, and a return to that same baseline roughly 3-4 s after sound offset (**Fig. 2B**).

The pupil-response evoked by the long auditory stimulus (4 s) in Experiment 1 was bi-phasic (**Fig. 2B**, top). The time-to-peak and height of the first response component was similar to what we observed in Experiment 2 for short auditory stimuli (100 ms): time-to-peak, ∼700ms; height, ∼3% signal change. The second response component (in Experiment 1) lasted throughout the 4s long auditory stimulus. The first transient may reflect the presence (possibly combined with unexpectedness) while the second sustained response component may reflect the characteristics (duration and white noise) of the task-irrelevant sound.

We quantified the size of the task-irrelevant sound-evoked pupil response as the mean of this differential time course within a time window of 2 s following task-irrelevant sound onset. We observed a positive effect of task-irrelevant sound on pupil size in virtually every participant (**Fig. 2C**). Repeated measures ANOVA (i.e., a t-test in case of Experiment 1) revealed a main effect for condition (no-task-irrelevant sound and task-irrelevant sound conditions) on pupil response in both experiments (Experiment 1: t_(21)_=14.665, p<0.001; Experiment 2: F_(8, 320)_=18.578, p<0.001). Bayesian t-tests additionally indicated that task-irrelevant sounds very likely caused an increase in pupil-linked arousal (Methods) in all timing windows or conditions of all experiments (so across different frequencies of task-irrelevant sound occurrence) given the respective data (BF_10_ range from 2.25e^7^ to 1.23e^11^). The magnitude of task-irrelevant sound-evoked pupil responses was relatively stable over the time course of the experimental blocks, showing no sign of habituation (**Fig. S2A**) and seemed to largely depend on the duration of the task-irrelevant sound (**Fig. 2B,C**, compare top to bottom).

In sum, auditory white noise stimuli evoked reliable pupil responses, which were superimposed onto the task-evoked responses and precisely timed.

### Amplitude of task-evoked pupil responses predicts reduction of choice bias

We sought to replicate the previously established (negative) correlation between task-evoked pupil-linked arousal and absolute bias in perceptual choice (de Gee et al., 2017; de Gee et al., 2020; Lewandowska et al., 2019; Schriver et al., 2020). In line with earlier work using the contrast detection task (de Gee et al., 2017; de Gee et al., 2020), participants exhibited a substantial and consistent conservative bias (group average signal detection theoretic criteria ± S.E.M.: 0.381 ± 0.072 and 0.491 ± 0.056 for Experiments 1 and 2, respectively; **Fig. S1B**).

As expected from the previous results (de Gee et al., 2017; de Gee et al., 2020; Lewandowska et al., 2019; Schriver et al., 2020), the magnitude of task-evoked pupil responses was negatively correlated with (absolute) bias (group average correlation coefficients r: Experiment 1, r=-0.211, t_(21)_=-3.953, p=0.001; Experiment 2, r=-0.297, t_(40)_=-4.645, p<0.001; **Fig. 3A**). We did not observe a consistent relationship between pupil responses and sensitivity (**Fig. S3A**) or reaction time (**Fig. S3C**), also in line with earlier work (de Gee et al., 2017; de Gee et al., 2020). Experiment 1 included two types of experimental blocks that differed in terms of target signal frequency (presence of gratings on 40% vs. 60% of trials). Accordingly (de Gee et al., 2020; Green & Swets, 1966), participants had a less conservative bias on the “frequent-signal” blocks than on the “rare-signal” blocks (**Fig. S4A**). However, there was no interaction between block type and the task-evoked pupil response effect on absolute bias (**Fig. S4B, C**).

**Figure 3.**
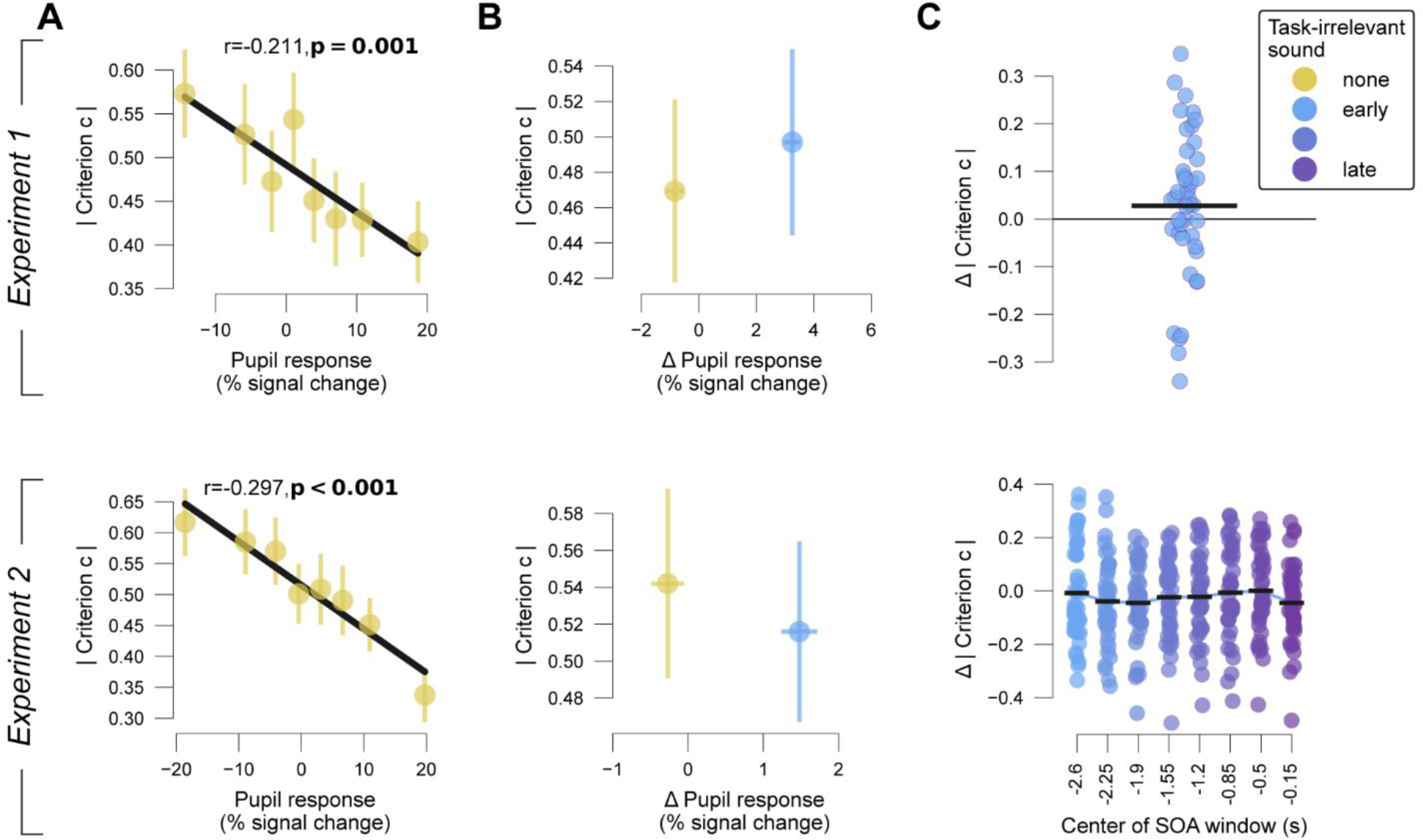
Pupil responses and choice bias. **(A)** Absolute choice bias (criterion c) plotted against task-evoked pupil response for experiments 1 and 2, for trials without a task-irrelevant sound. Error bars, S.E.M. across participants. **(B)** Mean absolute choice bias (criterion c) on trials with and without task-irrelevant sound plotted against mean task-irrelevant sound-evoked pupil responses. Error bars, S.E.M. across participants. **(C)** Participant-wise difference in absolute bias on task-irrelevant sound versus no task-irrelevant sound trials. Black bars, mean across participants.

### Task-irrelevant sound has no consistent effect on elementary decisions

Having established precise experimental control of pupil responses via the task-irrelevant sounds, we assessed their impact on decision-making. Task-irrelevant sounds did not affect choice bias (**Fig. 3C**, Experiment 1: t_(21)_=1.199, p=0.237; Experiment 2: F_(8, 320)_=1.239, p=0.276). The differences in absolute criterion between trials with and without task-irrelevant sound supported the null-hypothesis (BF_10_ ranging from 0.169 to 0.873). There was also no clear and consistent effect of the task-irrelevant sound on sensitivity or reaction time across the two experiments (**Fig. S5**) with one noteworthy exception: the task-irrelevant sound changed the speed-accuracy trade-off in Experiment 1 (but not Experiment 2). The task-irrelevant sound reduced reaction time (ΔRT=-0.086 ± 0.013 s, t_(21)_=-6.616, p<0.001, BF_10_=12370.926) and sensitivity (difference in d’=-0.126 ± 0.045 s.d., t_(21)_=-2.779, p=0.008, BF_10_=4.508). Intrinsic variability in task-irrelevant sound-evoked pupil-response did not correlate with absolute choice bias (**Fig. S6**). Taken together, task-irrelevant sounds had no consistent effect on choice behavior.

### Task-irrelevant sound does not alter weighing of evidence on choice

Our two previous experiments addressed simple perceptual decisions about the presence of a visual target signal superimposed on noise. Studies of more complex decisions requiring belief updating in the face of sequentially presented, discrete evidence samples reported that pupil responses predict the weight participants assign to new evidence while updating their beliefs (Filipowicz et al., 2020; Krishnamurthy et al., 2017; Murphy et al., 2021). To complete the picture of the effects of boosting phasic pupil-linked arousal with results in this more cognitively challenging perceptual decision-making domain, we wanted to know if boosting arousal by means of task-irrelevant sounds result in a similar upweighting of concomitant sensory evidence, or if, as for the null effect on overall choice bias, there is no such effect. To test this, we used a perceptual decision-making task with discrete evidence samples (Wyart et al., 2012), so that we could readily compute so-called “psychophysical kernels” that quantify the time course of the weighting of sequentially presented sensory evidence on the final behavioral choice (Okazawa et al., 2018; Waskom et al., 2019). Additionally, the placement of category boundaries implied a non-monotonic mapping from sensory (orientation) information to decision-relevant evidence (Wyart et al., 2012) and rendered the task more challenging than standard perceptual choice tasks.

In this Experiment 3 (**Fig. 4A**), participants reported whether the average orientation of a sequence of eight Gabor gratings of varying orientation was closer to the cardinal or diagonal axis (Wyart et al., 2012). The task-irrelevant sound started either at the same time as the baseline interval, together with the first grating, or together with the fifth grating in the sequence of eight, lasted for 200 ms and occurred at a pseudo-random 75% of trials. The design of Experiment 3 complicated the quantification of intrinsic variations of task-evoked pupil response: each trial contained a sequence of evidence samples of varying strengths, which could elicit arousal responses driven by uncertainty or surprise of different magnitudes within and across trials (Murphy et al., 2021; Urai et al., 2017). Thus, in Experiment 3 we did not characterize the intrinsic trial-to-trial variability in task-evoked pupil responses, as we did for Experiments 1 and 2 (**Fig. 2A**), but focused exclusively on the impact of task-irrelevant sound and the resulting pupil response.

**Figure 4.**
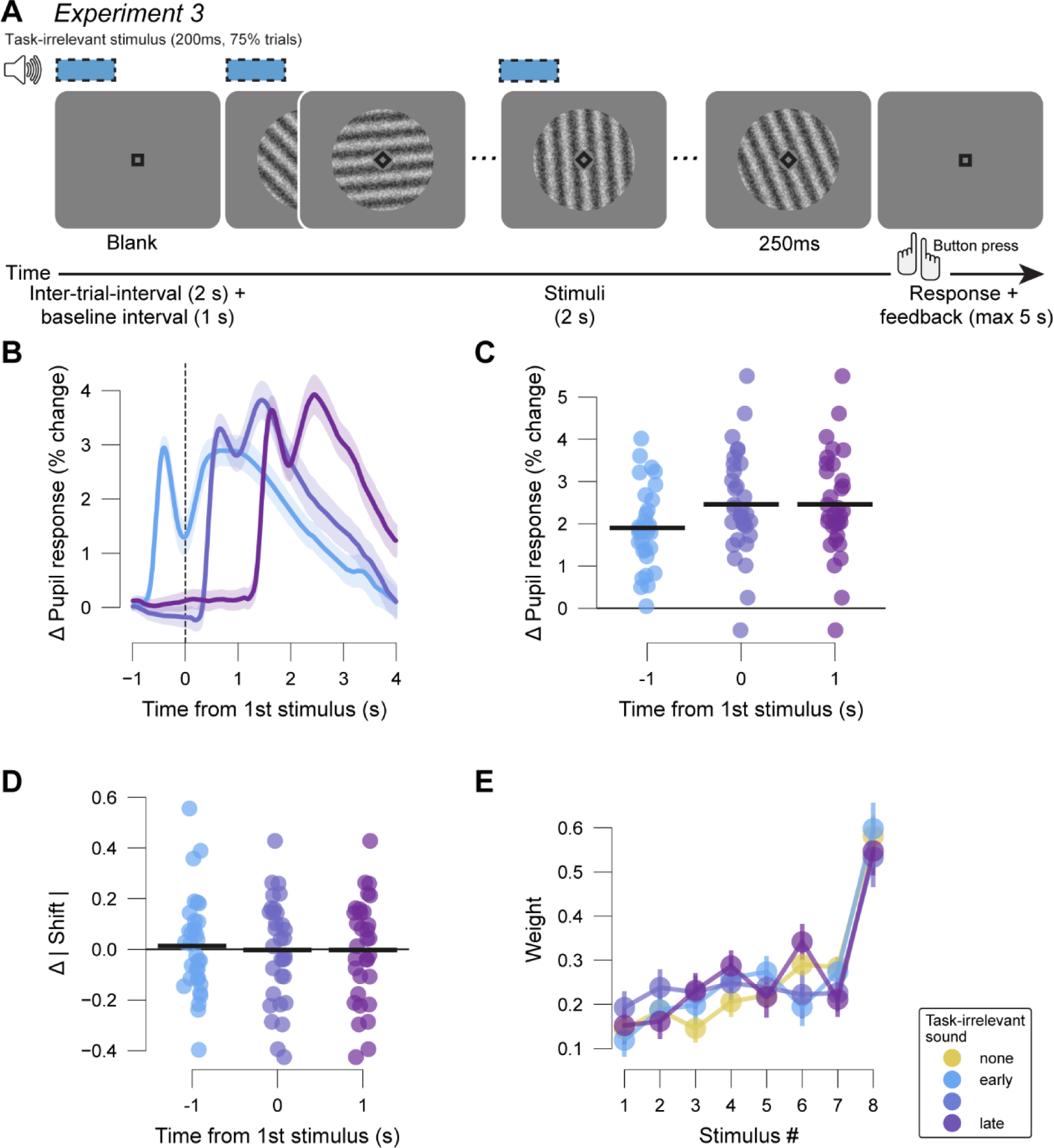
Task design, pupil responses and choice biases in Experiment 3. **(A)** Schematic sequence of events during the category level averaging task of Experiment 3. Participants reported the category (cardinal vs. diagonal) of the average orientation of gratings shown during each trial. White noise stimuli (75 dB) were played at the start of the inter-trial-interval, the first grating or the fifth grating (75% of trials). **(B)** Differential pupil size time courses between task-irrelevant sound trials and trials without task-irrelevant sound by task-irrelevant sound timing. Shading, S.E.M across participants. Baselined with 0.5 second intervals from before the first possible task-irrelevant sound. **(C)** Participant-wise task-irrelevant sound-evoked pupil response magnitude. Black bars, mean across participants. **(D)** Participant-wise difference in absolute bias (shift of psychometric function) on task-irrelevant sound versus no task-irrelevant sound trials. Black bars, mean across participants. **(E)** Regression weights of visual stimulus position as predictors of choice (psychometric kernels). Error bars, S.E.M across participants.

We quantified the pupil responses specifically evoked by the task-irrelevant auditory white noise stimuli in the same way as in Experiments 1 and 2. The mean resulting time courses showed robust increases in pupil size with respect to pre-trial baseline, and a return to that same baseline roughly 3-4 s after sound offset (**Fig. 4B**). A repeated measures ANOVAs revealed a main effect for condition (no-task-irrelevant sound and task-irrelevant sound conditions) on pupil response (Experiment 3: F_(3,93)_=71.742, p<0.001) (**Fig. 4C**). Bayes factor supported that task-irrelevant sounds very likely caused an increase in pupil-linked arousal in all timing conditions (BF_10_ range from 5.5e^9^ to 2.35e^11^).

Participants exhibited a substantial and consistent bias towards the diagonal choice category (group average shift of psychometric function ± S.E.M.: 0.36 ± 0.059). Task-irrelevant sounds, however, did not change absolute choice bias (**Fig. 4D**; F_(3, 93)_=0.313, p=0.816; BF_10_ for differences ranging from 0.189 to 0.232), nor sensitivity (slope of psychometric function; **Fig. S7B**; F_(3, 93)_=1.059, p=0.37; BF_10_ for differences ranging from 0.267 to 1.173).

We also estimated the weighting of evidence samples on choice using logistic regression (Methods). On trials without a task-irrelevant sound, participants exhibited an overall recency bias: a tendency to rely more on the most recent sample(s) of evidence (**Fig. 4E**). Critically, the psychophysical kernels on task-irrelevant sound trials were not different from the one computed on trials without a task-irrelevant sound (**Fig. 4E**). Taken together, task-irrelevant sounds also did not seem to alter the overall bias nor evidence weighing in this more complex task.

### No choice effect of task-irrelevant sound dependent on baseline pupil size

Having established an absence of behavioral effect of task-irrelevant sounds on different perceptual decision-making tasks, we finally tested for interaction of task-irrelevant sound effects on the pupil and choice bias with baseline pupil size. The amplitudes of task-evoked and task-irrelevant sound-evoked pupil responses were roughly stable across trials within blocks (**Fig. S2B, C**). By contrast, we observed large changes of the pre-trial baseline pupil sizes throughout the experiment, with a gradual constriction early in the block down to an asymptotic level (**Fig. S2A**). This was unlikely due to light adaptation, since participants sat in the same dark room for at least 5 minutes prior to the start of the experiment blocks, and because in one experiment (Experiment 2) with multiple longer (10 s duration) breaks between blocks of 100 trials, the pupil size dilated again in each break, resetting the process of monotonic constriction during the next block, despite constant illumination levels (**Fig. S2A,** middle).

As expected from previous work (de Gee et al., 2014; de Gee et al., 2020), pupil responses evoked by the task-irrelevant sound were negatively related to pre-trial baseline pupil size (p<0.001 on all experiments; **Fig. 5A**). Critically, however, differences in choice bias between task-irrelevant sound trials and those without did not depend on the pre-trial baseline size of the pupil (**Fig. 5B**). As pre-trial pupil sizes may contain spill-over from previous evoked pupil responses from the previous trial, we also assessed the baseline pupil diameter in the break intervals in Experiment 2. We found no effect of pupil size during these intervals on task-irrelevant sound effects on absolute bias (F_(3,120)_=0.33, p=0.804) or task-irrelevant sound-evoked pupil response on the following 100 trials (F_(3,120)_=0.234, p=0.873). There was also no interaction between effects of task-irrelevant sound timing window and pupil size during the break intervals on task-irrelevant sound effects on absolute bias (timing window: F_(3,120)_=0.358, p=0.784; interaction: F_(3,120)_=0.48, p=0.888) or task-irrelevant sound-evoked pupil response on the following 100 trials (timing window: F_(3,120)_=1.21, p=0.309; interaction: F_(3,120)_=1.615, p=0.109).

**Figure 5.**
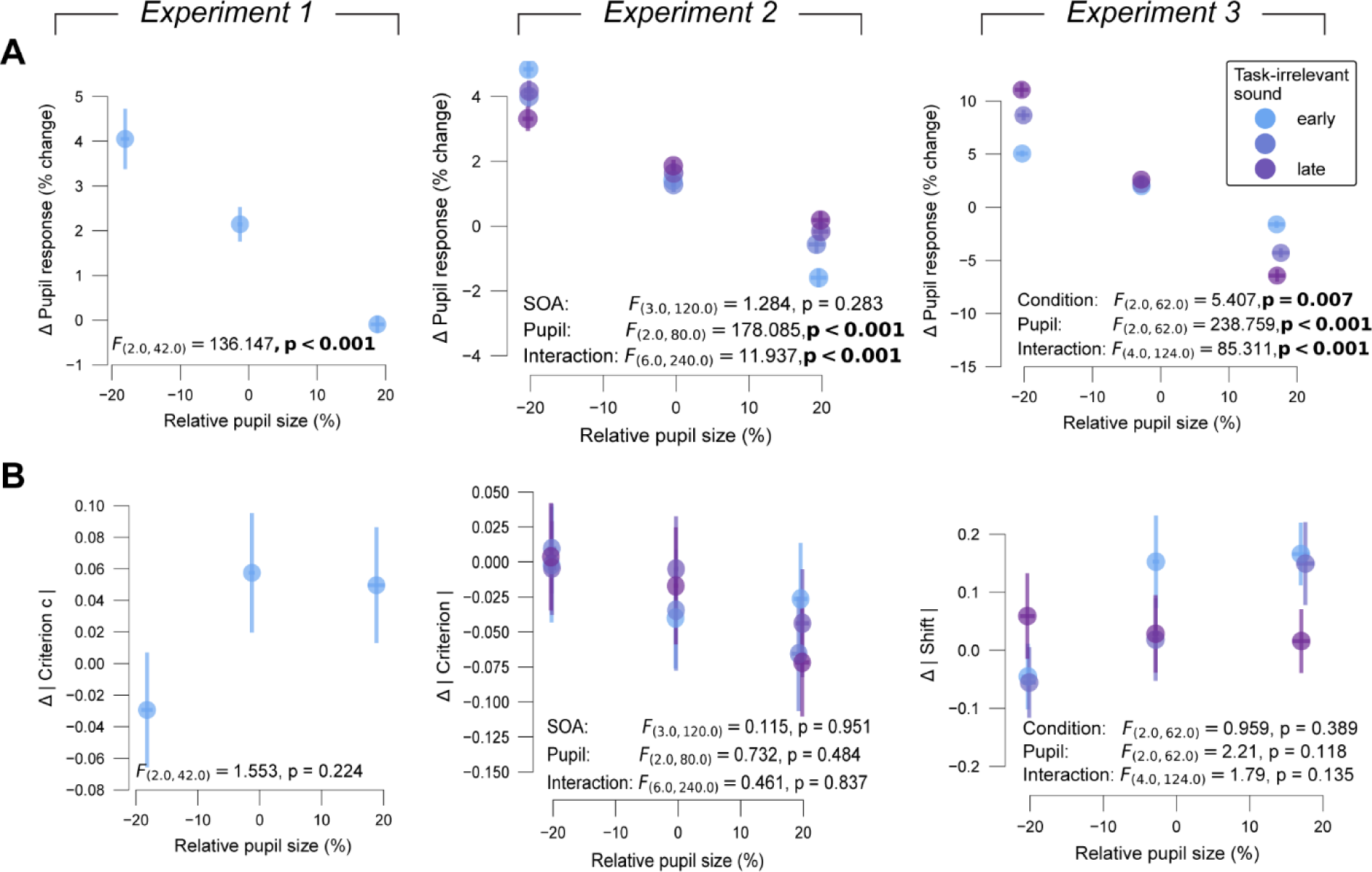
Effects of task-irrelevant sound as function of pre-trial baseline pupil size. **(A)** Task-irrelevant sound-evoked pupil response plotted against pupil baseline size (binned post-hoc by size). Shading, S.E.M. across participants. Statistical metrics, results of repeated measures ANOVA. **(B)** As (A) but for difference in absolute bias (criterion c, shift of psychometric function) between trials with and without task-irrelevant sound.

## Discussion

A common assumption underlying direct manipulations of brain circuits (Ross & Bassett, 2024) is that the experimentally elicited pattern of neural responses mimics the pattern of neural responses that the brain generates under physiological conditions. This assumption is typically implicit and rarely tested, especially in human subjects for which causal approaches to manipulate brain circuits in a specific fashion are limited to begin with. This caveat necessitates careful evaluation of any candidate approach for the experimental manipulation of neural events, including central arousal transients.

Here, within the same participants and experiments, we systematically compared the response profiles and behavioral correlates of phasic pupil-linked arousal responses that were task-related with those evoked by task-irrelevant sounds. We expected the latter to drive phasic responses of arousal-controlling brainstem nuclei, including the locus coeruleus (Joshi & Gold, 2020, 2022; Joshi et al., 2016). We conducted three experiments, in which participants performed challenging perceptual decisions while we varied the frequency, timing, and duration of the task-irrelevant sound. We replicated the negative correlation between intrinsic trial-to-trial variations in task-evoked pupil response amplitude and choice bias established in several previous studies (de Gee et al., 2017; de Gee et al., 2020; Krishnamurthy et al., 2017; Lewandowska et al., 2019; Schriver et al., 2020). As expected, the pupil did not only dilate during task engagement but also in response to the white noise sounds. All variants of this task-irrelevant sound drove robust and precisely timed pupil responses that were superimposed onto the task-evoked pupil responses and were separable through a simple linear subtraction. Yet, in neither experiment did these task-irrelevant sounds reduce choice bias, nor did we observe any other consistent behavioral effect.

The task-irrelevant sound-evoked pupil responses were smaller in amplitude than the range of intrinsic trial-by-trial variations of task-evoked pupil responses. Yet, the sound evoked-pupil responses were substantial in magnitude and reliable across subjects. Since the relationship between phasic pupil-linked arousal and choice bias is linear across the arousal’s full dynamic range (**Fig. 3A**), even a modest change in phasic arousal could, potentially, affect bias. Therefore, the smaller amplitude of sound-evoked compared to task-related pupil-linked arousal is unlikely to account for the complete absence of a behavioral correlate.

One previous study reported that task-irrelevant sounds can reduce choice bias (Bruel et al., 2022). However, in that task the auditory white noise was terminated upon the participant’s choice (button press). Thus, participants could control the offset of the slightly aversive sound stimulus, which may have caused them to favor speed over sensitivity, and which may have reduced choice bias. We took care to avoid this confound, and in three separate experiments did not find a consistent effect of task-irrelevant sounds on choice bias or other decision-making metrics.

Why did task-irrelevant sounds not influence decision-making despite producing reliable pupil responses? The interpretation depends on the assumption about the similarity of the central arousal events driven by the task and those driven by the task-irrelevant sound, which is currently unknown. If the two forms of central arousal events were identical (i.e., identical activation pattern of arousal-controlling brainstem systems), our result would imply that the correlation between trial-to-trial variations in task-evoked pupil responses and choice bias does not reflect a causal effect of arousal on behavior. For example, it is possible that forming a decision against one’s “default” choice option drives arousal. The latter could be conceptualized as a high-level form of surprise, akin to updating one’s internal belief with evidence that contradicts the prior belief, which drives pupil dilations (Filipowicz et al., 2020; Murphy et al., 2024; Murphy et al., 2021; Nassar et al., 2012; van den Brink et al., 2023). The time course of the correlation between task-evoked pupil responses and bias in reaction time versions of detection tasks shows that this relationship emerges long (∼800 ms) before a choice is reported (de Gee et al., 2017). Due to the delay of the pupil response relative to brainstem activity (Joshi et al., 2016), the underlying central arousal transient must have occurred even earlier. While it is possible that the content of evolving decisions *per se* changes central arousal state, it seems difficult to explain early pupil-behavior correlations by such (secondary) activation.

An alternative possibility is dissimilarity of the central arousal events recruited by task-irrelevant sounds and by the task. Evidence from fMRI suggests that pupil-linked bias reduction is mainly due to responses of neuromodulatory brainstem nuclei, rather than the superior and inferior colliculi (de Gee et al., 2017). Task-irrelevant sounds, on the other hand, seem to elicit less vigorous responses in the locus coeruleus than in the inferior colliculus (Joshi et al., 2016), which is a key stage of the central auditory pathway. Task-irrelevant stimuli might also recruit different sub-populations of neurons within the locus coeruleus than cognitive engagement. The locus coeruleus is more heterogeneously organized than long assumed (Poe et al., 2020; Totah et al., 2018). While the locus coeruleus is activated, to some extent, by both task engagement (Aston-Jones & Cohen, 2005) and sounds (Joshi et al., 2016), the relative amplitudes and neuronal sub-populations of these two types of locus coeruleus responses are unknown. Finally, multiple brainstem structures involved in the control of pupil size are strongly connected, providing a scaffold for complex interactions. For example, the locus coeruleus and the ventral tegmental area exhibit a close interplay (Ranjbar-Slamloo & Fazlali, 2019; Sara, 2009) and are co-activated with pupil responses during decisions (de Gee et al., 2017).

In sum, pupil responses recruited by task engagement (varying in amplitude from trial to trial) and driven by task-irrelevant stimuli may be determined by the relative contributions of several structures within the brain’s machinery controlling arousal state, with potentially different behavioral consequences. This idea relates to the conceptual distinction between arousing, alerting, and orienting (e.g. Berridge & Waterhouse, 2003; Poe et al., 2020; Wang & Munoz, 2015). For example, distinct functional processes have been postulated for executive, orienting, and alerting of attention (Posner & Boies, 1971; Raz & Buhle, 2006), whereby executive attention resembles the task recruitment we measured here, while alerting attention is a response to temporal cues, resembling our task-irrelevant sound.

While the pupil is likely to dilate in response to any sound, the level of unexpectedness of a task-irrelevant sound may be an important factor governing its behavioral effect. Indeed, in our experiments, the frequency of occurrence of the task-irrelevant sounds varied substantially (between 25% and 80%), and we found robust sound-evoked pupil responses throughout. Likewise, sounds that are completely predictable in terms of timing and content still drive pupil dilation (Knapen et al., 2016). This is expected simply because sounds elicit responses across the auditory pathways up to the inferior colliculus, which, in turn, can dilate the pupil (Joshi et al., 2016). By contrast, surprising events drive pupil responses in a more cognitive manner (de Gee et al., 2021; Filipowicz et al., 2020; Kloosterman et al., 2015; Lavín et al., 2013; Murphy et al., 2021; Preuschoff et al., 2011; van den Brink et al., 2023), including even responses to the absence of a sensory change, when change is expected (Filipowicz et al., 2020). This high-level component of the phasic arousal response may be critical for recruiting the locus coeruleus. It will be instructive to unravel the importance of these, and other characteristics (e.g., the nature of the sound) of task-irrelevant stimuli for producing central arousal transients that best mimic those driven by cognitive computation.

Previous work has reported that task-irrelevant stimuli produce a reduction in reaction time during cognitive tasks (Hackley et al., 2009; Hershenson, 1962; Jepma et al., 2009; Stahl & Rammsayer, 2005; Tona et al., 2016). Because of this convergent finding, the task-irrelevant stimuli in this literature were typically called “accessory stimuli”. We did not observe a consistent reduction of RT across all our current experiments. Only the least frequent but relatively long task-irrelevant sound in Experiment 1 (25% of trials; 4 s) caused speedier and less sensitive decision-making. The absence of an effect in Experiment 2 and 3 might be explained by the missing temporal binding, typically seen in the accessory stimulus literature. It may also be due to the relatively high frequency of trials on which an accessary stimulus occurred (80% and 75%, respectively), yielding the task-irrelevant sound to be a less surprising outcome. Yet another possibility is that our participants made challenging decisions that required protracted evidence accumulation, while the accessory stimulus literature mostly describes easier tasks that produced significantly shorter RTs (see Hackley et al., 2009; Hershenson, 1962; Jepma et al., 2009; Stahl & Rammsayer, 2005; Tona et al., 2016).

Across all experiments reported here, the pupil dilated reliably in response to the white noise bursts, regardless of timing, frequency, or time on task at which they were delivered. The mean value of the dilations seemed to scale with stimulus duration (although these durations were so far only varied across experiments that also differed in other variables) and the dilation latency precisely reflected the stimulus onset latency. This comprehensive assessment of pupil responses to task-irrelevant sounds complements and extends previous work in important ways (Cronin et al., 2023; Petersen et al., 2017; Tona et al., 2016). For example, Petersen et al. (2017) demonstrated the dependence of the pupil response amplitude on sound intensity, while we here demonstrated the dependence on sound duration and/or frequency, providing more room for future experimental manipulation (the permissible range of sound pressure levels in human experiments is quite limited). Most importantly, our work demonstrates that the pupil responses to task-irrelevant sounds can be readily isolated from the task-evoked responses using a simple linear subtraction approach as commonly used in fMRI studies. Like fMRI, this approach ignores possible non-linear interactions between the two types of superimposed arousal responses, which should be tested in future experiments. Yet, the clear stimulus dependence of the resulting estimates of the sound-evoked responses provides strong evidence for an approximately linear superposition.

We found that phasic pupil responses to our manipulation were reduced with increasing baseline pupil size. However, taking baseline pupil size into account did not change any results of decision-making behavior. Nassar et al. (2012) found effects of task-irrelevant sound-switches on learning rate that depended on the baseline pupil size. It is unclear whether our results can be compared to these findings as the behavior in question and the manipulations were different. Additionally, baseline pupil measures used for quantifying tonic arousal are often taken from short inter-trial-intervals and may thus be confounded by spillover of phasic pupil responses from previous trials (but see Beerendonk et al., 2024). We therefore chose to additionally use the purer average pupil size during small intervals of rest (10s) that appeared every 100 trials in Experiment 2. Pupil size during these intervals predicted neither behavioral nor pupil responses to the task-irrelevant sound.

To conclude, establishing approaches for non-invasively manipulating phasic arousal would in principle allow for causal tests of the role of central arousal transients in cognition and could be translated to practical applications in health and disease. For example, aberrancies in the central arousal system and in pupil responses are found in important neuropsychiatric disorders such as Parkinson’s Disease and Alzheimer’s Dementia (James et al., 2021; Jiménez et al., 2021; Tosserams et al., 2023). Here we critically evaluate one such candidate tool, task-irrelevant sounds presented during a challenging task. While we show that these sounds evoked one component of phasic arousal associated with pupil responses, the complexity of the central arousal system calls for careful consideration. This is in line with the emerging view that the brainstem system controlling central arousal state as well as pupil size is likely heterogenous, made up of different sub-systems with distinct functional roles. Future neuroimaging or direct recording studies should further illuminate and dissociate sources of these task-irrelevant stimulus-evoked and task-evoked arousal responses.

## Methods

### Participants

27 healthy participants (13 female; age range, 8-28 y) participated in Experiment 1. 41 healthy participants (29 female; age range, 18-35 y) participated in Experiment 2. 32 healthy participants (19 female; age range, 18-33 y) participated in Experiment 3. All participants had normal or corrected-to-normal vision. They gave written informed consent and were remunerated by the hour (Experiment 2) or received credit points (Experiments 1 and 3). Experiments 1 and 3 were approved by the ethics committee of the Department of Psychology at the University of Amsterdam, and Experiment 2 by the ethics committee of the Faculty of Psychology and Human Movement Sciences at the University of Hamburg. Five participants were excluded from Experiment 1 due to low number of trials (<1000) after removing trials with more than 20 % missing pupil data (see analysis of pupil responses) or reaction times below 200 ms or longer than 4.5 s leading to a total sample of 97 participants.

### Behavioral tasks

All participants were asked to sit in front of the screen at 50cm distance resting their head on a chin rest. They indicated their choice with a button press: ‘z’ or ‘m’ on a keyboard, counterbalanced across participants.

### Contrast detection task (Experiments 1 & 2)

Each trial of the contrast detection task (see de Gee et al., 2014) consisted of an inter-trial interval (ITI), a baseline interval, and a decision interval. The ITI showed a blank grey screen and a fixation square. It lasted 3-4 s in Experiment 1 (uniformly distributed) and 3.8 s in Experiment 2. The baseline interval (Experiment 1, 1 s; Experiment 2, 0.2-1.4 s, exponentially distributed) additionally showed flickering visual noise (refresh rate: 100Hz) in a gaussian annulus around the fixation square with 20% contrast. The decision interval was cued by a 45° rotation of the fixation square, and, on a fraction of trials showed a signal (Gabor grating; 2 cycles per degree; vertical orientation) superimposed onto the visual noise. In Experiment 1, in ‘rare blocks’, the signal appeared in 40% of trials, and in ‘frequent blocks’ on 60% of trials. In Experiment 2, the signal appeared in 50% of trials. Participants reported the presence or absence of the signal with a button press which ended the trial. The maximum possible duration of the decision interval was 5 s in Experiment 1 and 7 s in Experiment 2. Luminance levels were kept stable throughout the whole task. In Experiment 1, visual stimuli were displayed on a gamma-corrected monitor (spatial resolution of 2560 by 1440 pixels) with a vertical refresh rate of 100 Hz, and in Experiment 2 on a VIEWPixx monitor (1920 by 1080 pixels) with a refresh rate of 100 Hz.

A task-irrelevant sound (auditory white noise) occurred at a pseudorandom fraction of trials: Experiment 1, 25%; Experiment 2, 80%. In Experiment 1, the task-irrelevant sound (70 dB) started together with the baseline interval and lasted 4 s. In Experiment 2, the task-irrelevant sound (70 dB) was played randomly between 3 s pre and 0.5 s post decision interval onset (uniformly distributed) and lasted for 100 ms. In Experiment 1, task-irrelevant sounds were played from Logitech speakers, and in Experiment 2 through AKG K72 headphones.

Experiment 1 was conducted at the University of Amsterdam over three separate sessions. In the first session (∼ 1 h), participants were acquainted with the contrast detection task. We titrated individual task difficulty to about 75% accuracy by adjusting contrast levels of the target stimulus. In each of the sessions two and three (∼ 2.5 h each), participants underwent two blocks of the experiment (40 minutes, 360 trials per block) during which their EEG was recorded, and their eyes were tracked.

Experiment 2 was conducted at the University Medical Center Hamburg-Eppendorf. Participants came into the lab on two days within a week for two hours each day. The first day started with a contrast orientation discrimination task. In this task, the stimulus contrast changed following a staircase procedure aimed at 75% accurate performance. The last contrast level was used as first contrast level of the signal in the contrast detection task. Participants were then asked to perform 5-minute practice blocks of the contrast detection to get acquainted with it and to allow for further manual adjustment of contrast. After four to seven practice blocks, participants took a break and then continued with the first experiment block. On the second day, participants first performed another practice block and then performed two experiment blocks separated by a 5-minute break. Experiment blocks consisted of 400 trials, took approximately 45 minutes, and included eye tracking.

### Orientation averaging task (Experiment 3)

Each trial of the category level orientation averaging task (Wyart et al., 2012) consisted of an ITI, a baseline interval, a grating sequence interval, and a decision interval. The ITI showed a blank grey screen and a fixation square. It lasted 2 s. The baseline interval was identical and lasted 1 s. During the grating sequence interval, 8 Gabor gratings of individual orientation were shown consecutively for 250 ms each. The decision interval started with the end of the last grating and showed a grey screen with the fixation square. Participants reported whether the average orientation of the gratings belonged to the cardinal or diagonal category (true distribution 50%/50%) with a button press. A pair of tones played 250 ms after their button press served as feedback which ended the trial (correct: 440 followed by 880 Hz, incorrect: 880 followed by 440 Hz). Difficulty level was kept around 75% accuracy with the help of an online staircase procedure adjusting the distance between the angle of grating orientation and the respective perfect cardinal or diagonal angle. Luminance levels were kept stable throughout the whole task. Visual stimuli were displayed on a gamma-corrected monitor (spatial resolution of 2560 by 1440 pixels) with a vertical refresh rate of 100 Hz.

The task-irrelevant sound (auditory white noise) occurred at a pseudorandom 75% of trials in this experiment. It was 75 dB loud, 200 ms long, and started either at the onset of the baseline interval, simultaneously to the first Gabor grating or at the same time as the fifth Gabor grating (25% of trials each). Task-irrelevant sounds were played from Logitech speakers.

Experiment 3 was conducted at the University of Amsterdam in two sessions (∼ 2 hours each). In session one, participants completed two practice blocks (∼ 10 minutes each) before running three experiment blocks (∼ 25 minutes each; 200 trials). In session two, four more experiment blocks took place.

### Eye data acquisition

Eye data were obtained with Eyelink 1000 devices (SR Research, Osgoode, Ontario, Canada) at 1000 Hz with an average spatial resolution of 15 to 30 min arc. The eye trackers were calibrated at the start of each experiment block. In experiments 1 and 3 we recorded the left eye, and in Experiment 2 we recorded the right eye.

### Analysis of pupil responses

#### Preprocessing

Pupil data were analyzed with custom-made Python scripts like those used in previous studies (de Gee et al., 2017; de Gee et al., 2014). Blinks and saccades were detected by the manufacturer’s standard algorithms. Missed blinks were detected with a custom algorithm (van den Brink et al., 2016). Pupil data were low-pass filtered with a third-order Butterworth filter with a cut-off at 10Hz to correct for slow drift. Missing data (e.g. caused by blinks) were linearly interpolated in a window of 200 ms before up to 200 ms after the missing data. Pupil responses to blinks and saccades were corrected using a double gamma function convolution (Knapen et al., 2016). Using the median, the units of the pupil size time series were transformed from pixels to percent signal change.

#### Quantification of task-evoked pupil responses

To measure trial-wise task-evoked pupil responses, we created epochs centered on choice (button-press) for trials with no task-irrelevant sound. Epochs with more than 20% missing data (due to blinks or general signal loss) were excluded from the analyses. The epoched pupil size data was baselined with the mean of half a second before the first visual target stimulus could appear (decision interval onset). For each trial, we then calculated the mean (baseline subtracted) pupil size over two seconds starting 0.5 s before choice (button-press) (de Gee et al., 2014). To specifically isolate trial-to-trial variations of underlying response amplitudes, variations due to baseline pupil size and reaction time were removed from the task-evoked pupil responses (via linear regression; de Gee et al., 2017). In one analysis, trials were sorted by task-evoked pupil response amplitude and collapsed into eight bins (**Fig. 2A**).

#### Quantification of task-irrelevant sound-evoked pupil responses

For measuring the pupil responses evoked by the task-irrelevant sound in Experiment 1 and 3, we created epochs centered on task-irrelevant sound onset. Epochs with more than 20% missing data (due to blinks or general signal loss) were excluded from the analyses. Epochs were baselined using the mean pupil size from half a second before task-irrelevant sound onset. From each epoch, we then subtracted the block-wise average pupil time series on the trials without a task-irrelevant sound. For experiment 2, we used epochs centered on decision-interval onset. These were baselined using the mean pupil size from half a second before the first possible onset of task-irrelevant sounds. From these epochs, we subtracted the block-wise average pupil time series on the trials without a task-irrelevant sound. Epochs were then aligned to task-irrelevant sound onset and baselined again with the mean pupil size from half a second before task-irrelevant sound onset. For all experiments, we then calculated the mean (baseline subtracted) differential pupil size over two seconds following task-irrelevant sound onset for each trial.

#### Quantification of baseline pupil size

Pre-trial baseline pupil size was quantified as the mean pupil size (percent signal change) over 0.5 seconds before the first possible event in each trial. Experiment 2 included five 10 s-intervals in each block (every 100 trials). From these we extracted longer baseline values by averaging pupil size over 7 s starting 2 s after interval onset.

### Analysis of choice behavior

For all analyses, we excluded the first 10 trials of each block due to habituation effects (see **Fig. S2**).

In experiments 1 and 2, we computed signal detection theory metrics sensitivity d’ and criterion c (Green & Swets, 1966; Macmillan & Creelman, 2005). The sensitivity describes the participants’ ability to discriminate signal from noise and is calculated as the difference between the z-scored hit rate and false alarm rate. The choice bias describes the participants’ intrinsic tendency to prefer one choice alternative over the other and is calculated as the average of z-scored hit and false alarm rates multiplied by −1.

In Experiment 3, we used the slope and shift from a fitted psychometric function. The psychometric function was obtained from a logistic regression that used the normalized sample-wise (signed) evidence strength (i.e. orientation of each of the trial’s eight Gabor gratings) to predict choice. The average regression coefficient of the Gabor gratings quantified the slope reflecting sensitivity. The regression intercept quantified the shift reflecting choice bias. Similarly, psychophysical kernels were quantified as regression coefficients of sample-wise (signed) evidence strength (i.e. orientation; not normalized).

In Experiment 1 and 2, reaction time was defined as the time from decision interval onset until the button press. For analysis, these values were collapsed into 3 bins. In Experiment 3, reaction time was defined as the time from offset of the final stimulus in the sequence until the button press. Trials with reaction times lower than 0.2 s or higher than 4.5 s were excluded from analyses.

### Statistical comparisons

For each participant in Experiments 1 and 2, we computed the Pearson correlation between each of the behavioral metrics and the bin-wise average task-evoked pupil response. The sample of correlation coefficients was then compared against 0 with a paired-samples t-test. For Experiment 1, all metrics were calculated by block type and then averaged (**Fig. 2C**, **3C**) or tested separately (**Fig. S4**).

We compared pupil size and behavioral metrics on trials with a task-irrelevant sound versus trials without. In Experiment 2, we binned trials with a sliding window along the possible task-irrelevant sound onset times (length, 800 ms; step size, 350 ms). In Experiment 3, we considered the three possible task-irrelevant sound onset times separately. We assessed the task-irrelevant sound effect with repeated measures ANOVAs with (no-)task-irrelevant sound condition (or window) as within-subject factor. Further, differences between each task-irrelevant sound condition or time window and the no task-irrelevant sound condition was tested against zero using Bayesian t-tests (**Fig. 2C, 3C, 4C,D, S5**).

For analysis of the psychophysical kernels in Experiment 3, we tested each kernel of each condition with a task-irrelevant sound against the respective kernel of the condition without a task-irrelevant sound using FDR-corrected Wilcoxon tests (**Fig. 4E**).

We checked for interaction of task-irrelevant sound effects behavior and pupil response with pre-trial baseline pupil size by means of repeated measures ANOVA. We tested the effects of the within-subject factor pre-trial pupil baseline bin on task-irrelevant sound-evoked pupil response or on difference in absolute bias towards the condition without task-irrelevant sounds (**Fig. 5**). For Experiments 1 and 3, this ANOVA also included the factor task-irrelevant sound condition (i.e., task-irrelevant sound SOA time windows in Experiment 2). Additionally for Experiment 2, we calculated an ANOVA testing the effects of the 10-s break interval pupil size (as well as task-irrelevant sound SOA window) on the effects of the task-irrelevant sound on absolute bias or pupil response in the following 100 trials. For this analysis, task-irrelevant sound SOAs were binned into 4 bins with cut-offs at −2.125 s, −1.25s, and −.375 s with regard to decision interval onset.

## Data and code availability

Data and analysis scripts will be made publicly available upon publication.

## Funding

This work was funded by the Deutsche Forschungsgemeinschaft (DFG, German Research Foundation) - GRK 2753/1 - Project number 449640848 (to LS, THD)

## Acknowledgements

We thank Noé Igounenc and Joffrey Straub for help with data collection in Amsterdam.

## Author Contributions

Conceptualization: JH, LS, THD, JWdG

Experimental design: JH, ACG, EZ, MB, LS, SvG, THD, JWdG

Data acquisition: JH, ACG, EZ, MB

Formal analysis: JH, JWdG

Writing—original draft: JH, THD, JWdG

Writing—review and editing: JH, SvG, LS, THD, JWdG

Supervision: THD, JWdG

Funding acquisition: LS, THD

## Competing Interests

The authors declare no competing interests.

## Supplementary figures

**Figure S1.**
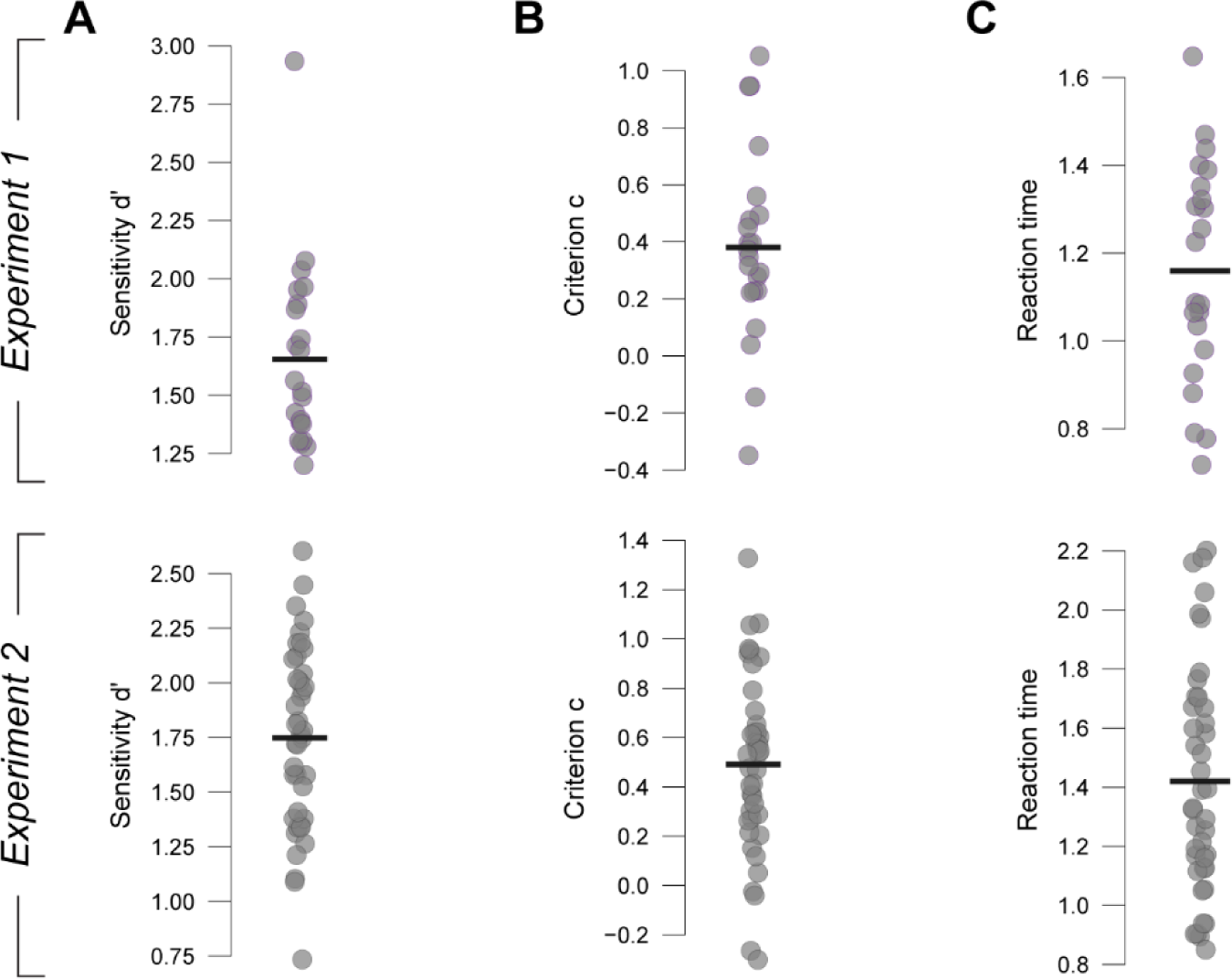
Mean behavior. **(A)** Mean individual decision performance values (sensitivity d’, slope of psychometric function) for Experiments 1 (mean ± S.E.M: 1.655 ± 0.085) and 2 (1.748 ± 0.065). Black bar, group mean. **(B, C)** As A, but for choice bias (mean criterion c ± S.E.M.: Experiment 1, 0.381 ± 0.072; Experiment 2, 0.491 ± 0.056; B) and reaction time (mean ± S.E.M.: Experiment 1, 1.16 ± 0.054 s; Experiment 2, 1.42 ± 0.06 s; C).

**Figure S2.**
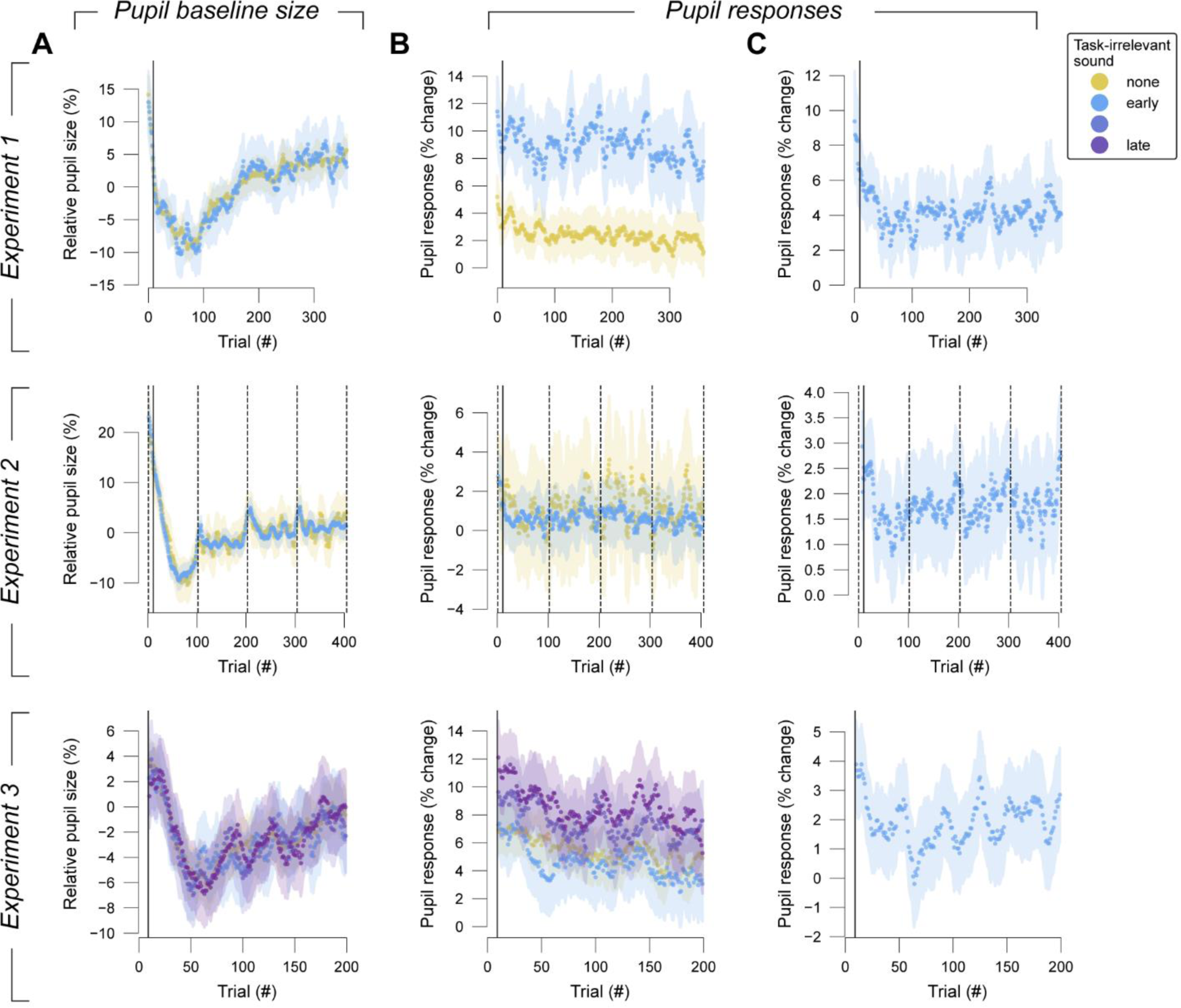
Temporal evolution of baseline pupil sizes, task-irrelevant sound-evoked pupil responses, and task-evoked pupil responses across the experiments. All three measures are shown as a sequence of group-average scalar amplitude values per trial. **(A)** Pre-trial pupil baseline per trials on all blocks by experiment, expressed as percent modulation around the median pupil size across the complete recording. Shaded area, S.E.M. across participants. Line, trial cut-off used for analyses. Dashed line, 10 s-break interval in Experiment 2. **(B, C)** As A, but for task-evoked pupil response (B) and task-irrelevant sound-evoked pupil response (C). Pupil response amplitudes are expressed as percent change relative to pre-trial baseline.

**Figure S3.**
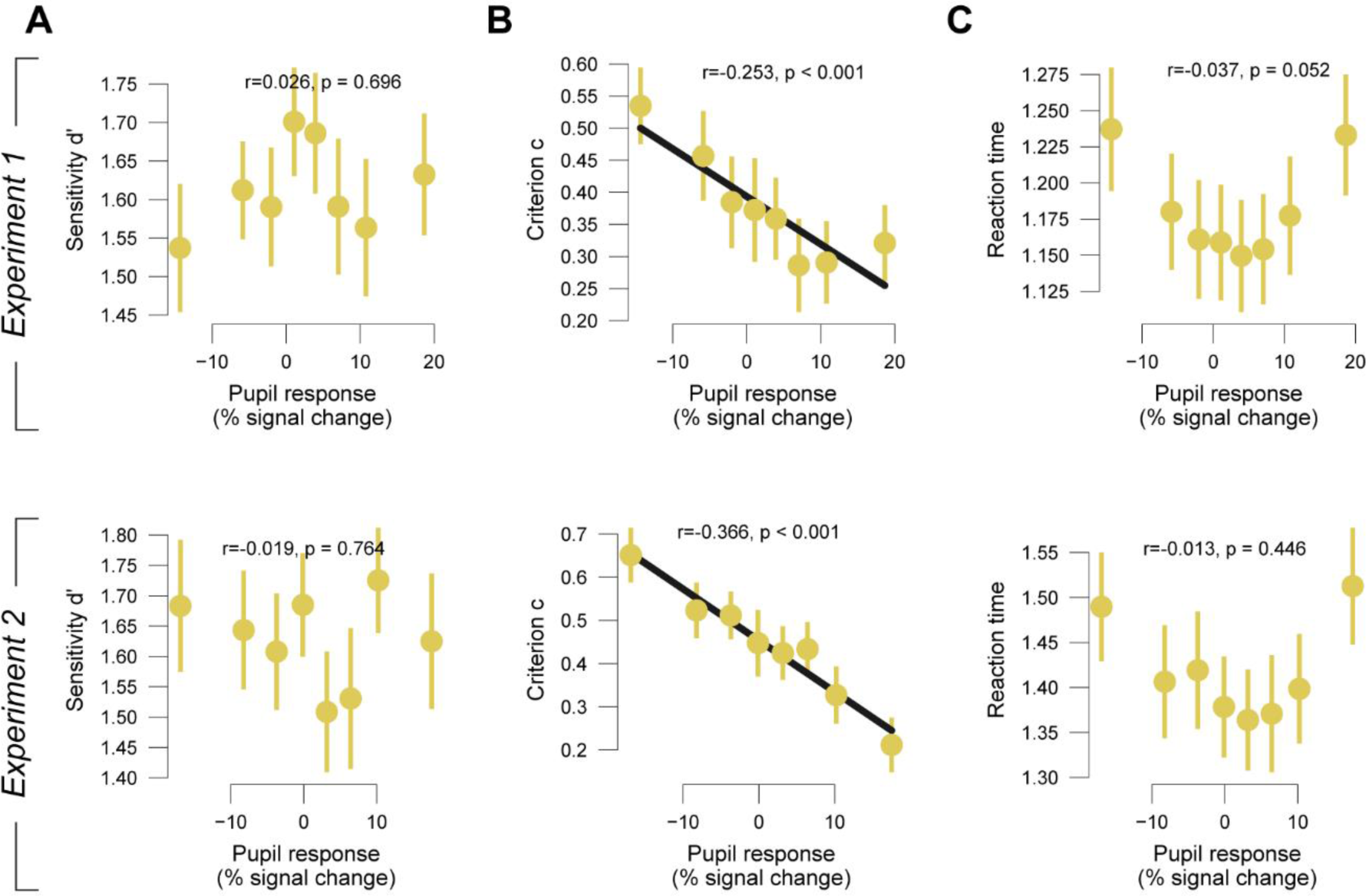
Correlation analyses between TPR and behavior. **(A)** Decision sensitivity d’ plotted against task-evoked pupil response for experiments 1 and 2. Experiment 3 was not included in this analysis. Error bars, S.E.M. across participants. Only trials with no sound were used for these analyses. **(B, C)** As A, but for choice bias (criterion c; B) and reaction time (C).

**Figure S4.**
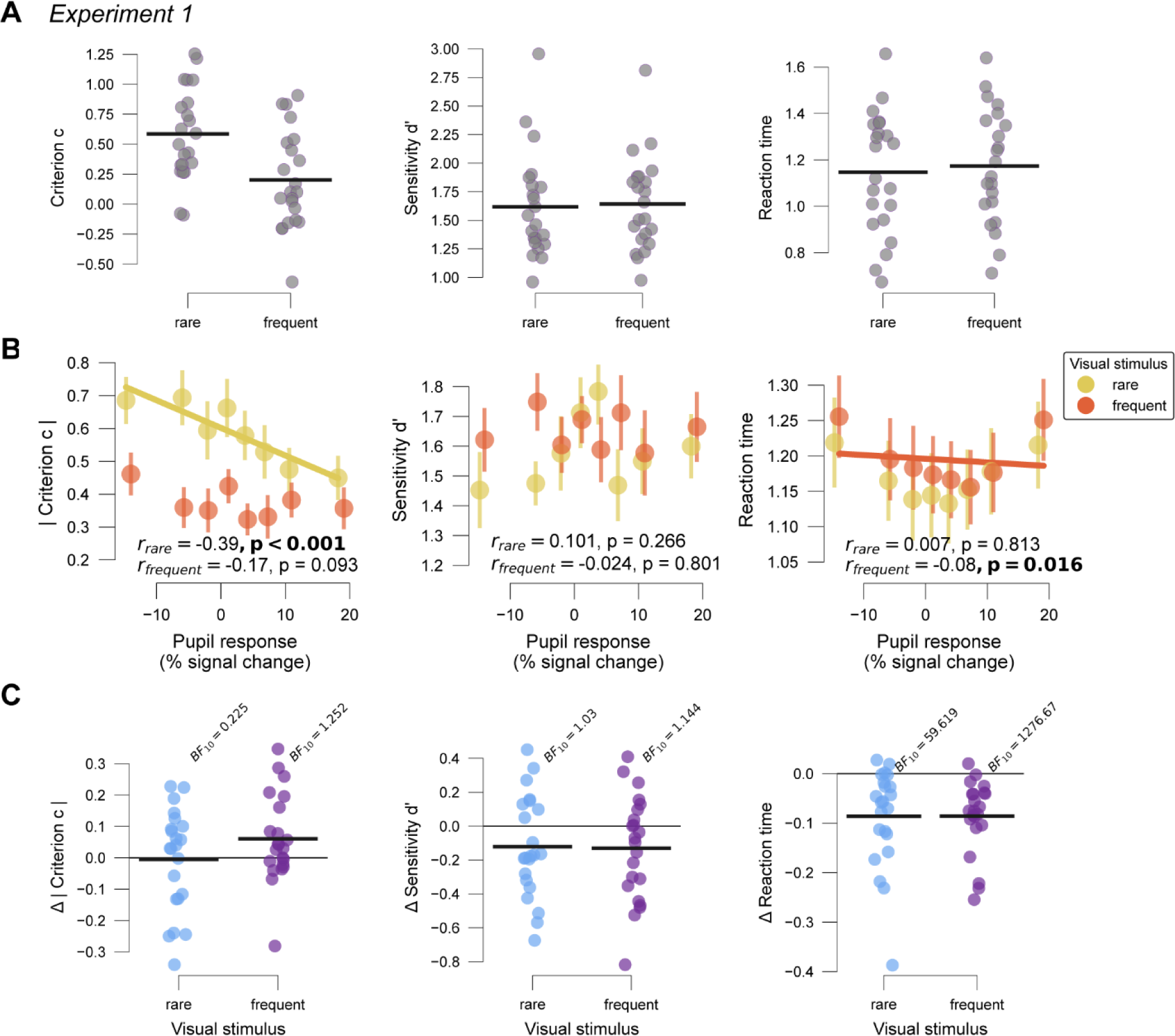
Experiment 1 Behavioral effects during task-evoked pupil responses and of task-irrelevant sound by block type. **(A)** Mean individual decision bias (criterion c), sensitivity (d’) and reaction time values by block type. Black bar, group mean. **(B)** Absolute criterion c, sensitivity d’ and reaction time plotted against task-evoked pupil response. Colors represent block type. Error bars, S.E.M. across participants. Only trials without task-irrelevant sound were used for these analyses. **(C)** Individual mean differences in absolute criterion c, sensitivity d’ and reaction time between trials with and without task-irrelevant sound. Black bar, mean. BF_10_ values for Bayesian t-tests. Colors represent block type.

**Figure S5.**
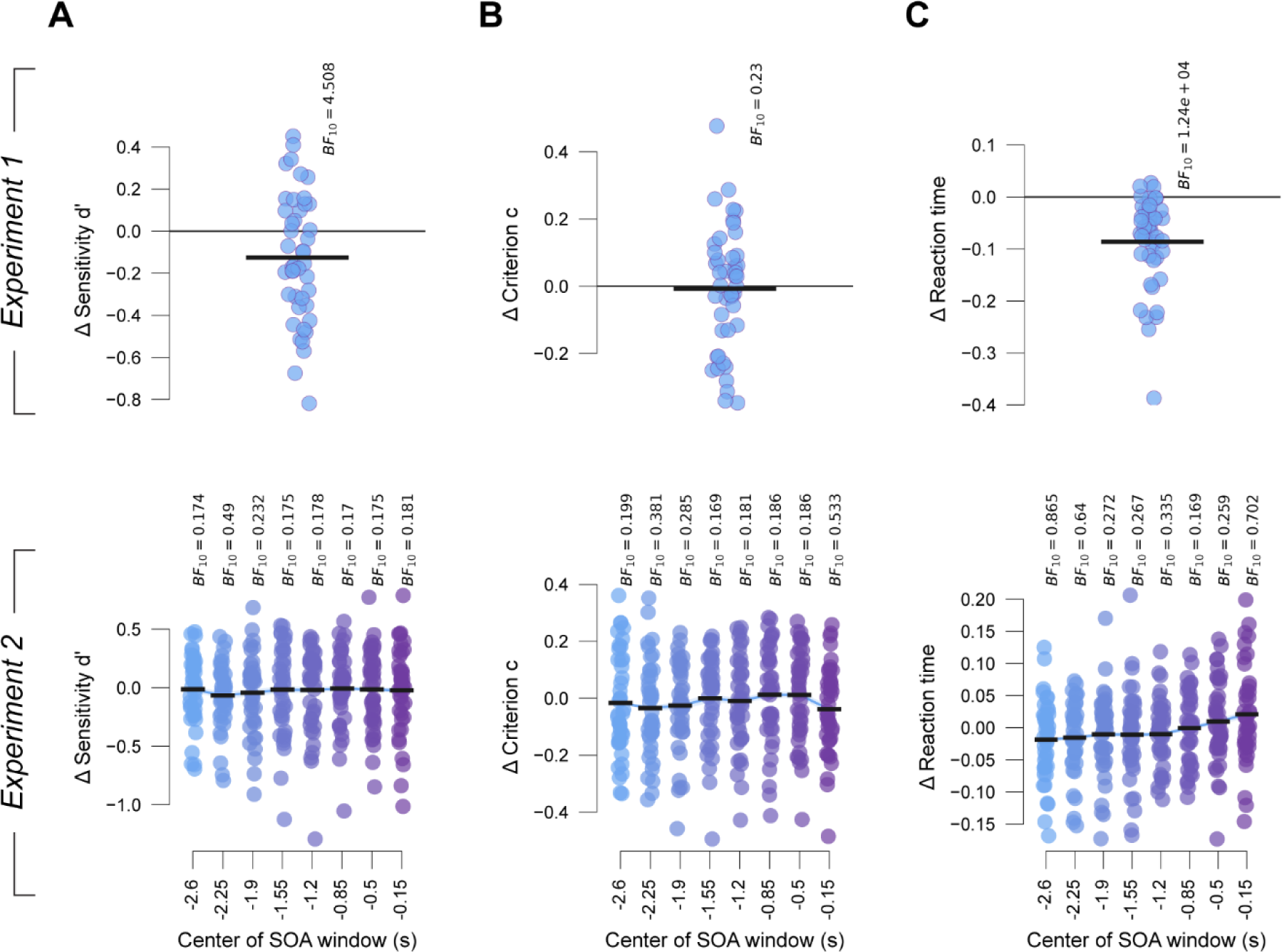
Behavioral responses to task-irrelevant sound. **(A)** Individual mean differences in sensitivity (d’) between trials with and without task-irrelevant sound. Black bar, mean. BF_10_ values for Bayesian t-tests. **(B, C)** As A, but for bias (criterion c) and reaction time, respectively.

**Figure S6.**
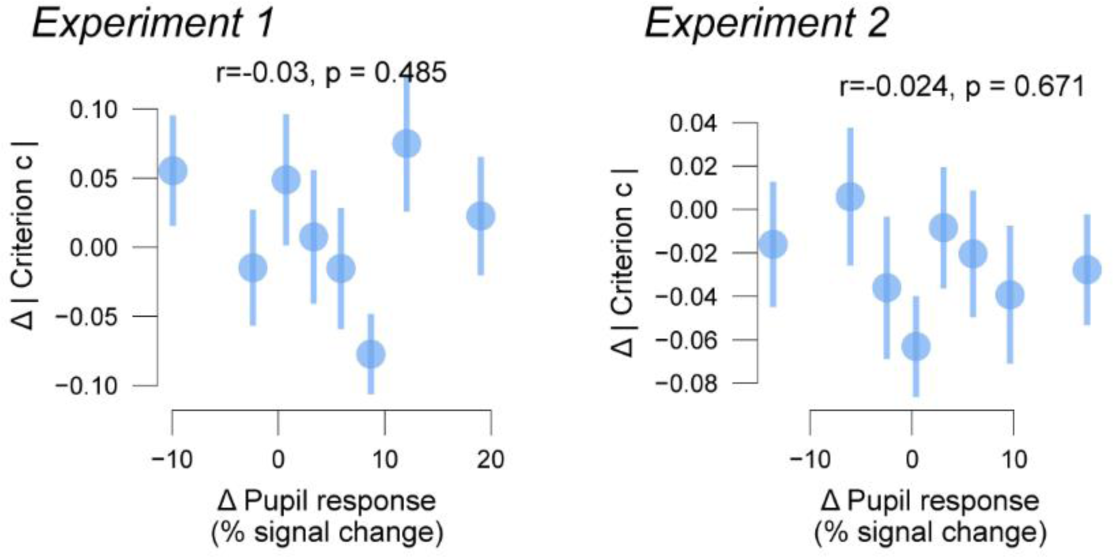
Variability of task-irrelevant sound-evoked pupil response. Absolute bias (criterion c) plotted against pupil response to the task-irrelevant sound. Error bars, S.E.M. across participants.

**Figure S7.**
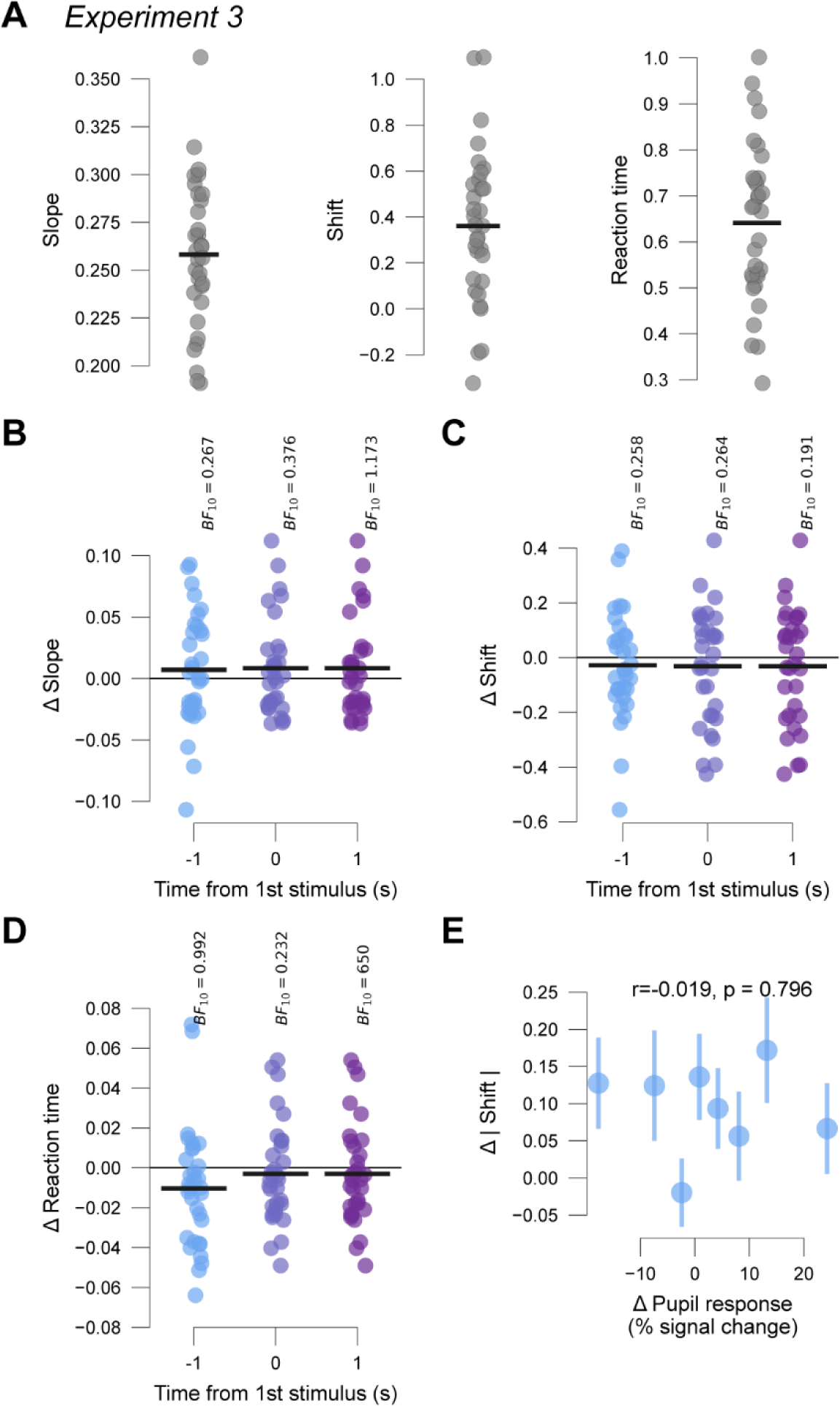
Experiment 3 behavioral means and task-irrelevant sound-evoked effects. **(A)** Mean participant-wise performance (mean slope of psychometric function ± S.E.M. across participants: 0.258 ± 0.007), choice bias (mean shift of psychometric function ± S.E.M. across participants: 0.36 ± 0.059) and reaction time (mean ± S.E.M. across participants: 0.642 ± 0.031 s). Black bars, overall mean across participants. **(B)** Individual mean differences in performance between trials with and without task-irrelevant sound. Black bar, mean. BF_10_ values for Bayesian t-tests. Colors represent different task-irrelevant sound timing conditions. **(C, D)** As B, but for choice bias and reaction time, respectively. **(E)** Absolute bias plotted against pupil response to the task-irrelevant sound. Error bars, S.E.M. across participants.

## Notes

### Competing Interest Statement

The authors have declared no competing interest.

## References

Arnsten, A. F. T. (2015). Stress weakens prefrontal networks: Molecular insults to higher cognition. Nature Neuroscience, 18(10), 1376–1385. 10.1038/nn.4087

Aston-Jones, G., & Cohen, J. D. (2005). An integrative theory of locus coeruleus-norepinephrine function: Adaptive gain and optimal performance. Annual Review of Neuroscience, 28, 403–450. 10.1146/annurev.neuro.28.061604.135709

Beatty, J. (1982). Task-evoked pupillary responses, processing load, and the structure of processing resources. Psychological Bulletin, 91(2), 276–292. 10.1037//0033-2909.91.2.276

Beerendonk, L., Mejías, J. F., Nuiten, S. A., de Gee, J. W., Fahrenfort, J. J., & van Gaal, S. (2024). A disinhibitory circuit mechanism explains a general principle of peak performance during mid-level arousal. Proceedings of the National Academy of Sciences, 121(5), e2312898121. 10.1073/pnas.2312898121

Berridge, C. W., & Waterhouse, B. D. (2003). The locus coeruleus-noradrenergic system: Modulation of behavioral state and state-dependent cognitive processes. Brain Research Reviews, 42(1), 33–84. 10.1016/S0165-0173(03)00143-7

Bouret, S., & Sara, S. J. (2005). Network reset: A simplified overarching theory of locus coeruleus noradrenaline function. Trends in Neurosciences, 28(11), 574–582. 10.1016/j.tins.2005.09.002

Breton-Provencher, V., Drummond, G. T., Feng, J., Li, Y., & Sur, M. (2022). Spatiotemporal dynamics of noradrenaline during learned behaviour. Nature, 606(7915), 732–738. 10.1038/s41586-022-04782-2

Breton-Provencher, V., & Sur, M. (2019). Active control of arousal by a locus coeruleus GABAergic circuit. Nature Neuroscience, 22(2), 218–228. 10.1038/s41593-018-0305-z

Bruel, B. M., Katopodis, V. G., de Vries, R., Donner, T. H., McGinley, M. J., & de Gee, J. W. (2022). Auditory accessory stimulus boosts pupil-linked arousal and reduces choice bias. BioRxiv, 2022-08. 10.1101/2022.08.28.505585

Cheadle, S., Wyart, V., Tsetsos, K., Myers, N., de Gardelle, V., Herce Castañón, S., & Summerfield, C. (2014). Adaptive gain control during human perceptual choice. Neuron, 81(6), 1429–1441. 10.1016/j.neuron.2014.01.020

Cronin, S. L., Lipp, O. V., & Marinovic, W. (2023). Pupil dilation during encoding, but not type of auditory stimulation, predicts recognition success in face memory. Biological Psychology, 178, 108547. 10.1016/j.biopsycho.2023.108547

Dayan, P., & Yu, A. J. (2006). Phasic norepinephrine: A neural interrupt signal for unexpected events. Network: Computation in Neural Systems, 17(4), 335–350. 10.1080/09548980601004024

de Gee, J. W., Colizoli, O., Kloosterman, N. A., Knapen, T., Nieuwenhuis, S., & Donner, T. H. (2017). Dynamic modulation of decision biases by brainstem arousal systems. ELife, 6, e23232. 10.7554/eLife.23232.001

de Gee, J. W., Correa, C. M. C., Weaver, M., Donner, T. H., & van Gaal, S. (2021). Pupil Dilation and the Slow Wave ERP Reflect Surprise about Choice Outcome Resulting from Intrinsic Variability in Decision Confidence. Cerebral Cortex, 31(7), 3565–3578. 10.1093/cercor/bhab032

de Gee, J. W., Knapen, T., & Donner, T. H. (2014). Decision-related pupil dilation reflects upcoming choice and individual bias. Proceedings of the National Academy of Sciences of the United States of America, 111(5), E618–25. 10.1073/pnas.1317557111

de Gee, J. W., Tsetsos, K., Schwabe, L., Urai, A. E., McCormick, D., McGinley, M. J., & Donner, T. H. (2020). Pupil-linked phasic arousal predicts a reduction of choice bias across species and decision domains. ELife, 9. 10.7554/eLife.54014

Filipowicz, A. L., Glaze, C. M., Kable, J. W., & Gold, J. I. (2020). Pupil diameter encodes the idiosyncratic, cognitive complexity of belief updating. ELife, 9. 10.7554/eLife.57872

Gilzenrat, M. S., Nieuwenhuis, S., Jepma, M., & Cohen, J. D. (2010). Pupil diameter tracks changes in control state predicted by the adaptive gain theory of locus coeruleus function. Cognitive, Affective, & Behavioral Neuroscience, 10(2), 252–269. 10.3758/CABN.10.2.252

Grant, S. J., Aston-Jones, G., & Redmond, D. E. (1988). Responses of primate locus coeruleus neurons to simple and complex sensory stimuli. Brain Research Bulletin, 21(3), 401–410. 10.1016/0361-9230(88)90152-9

Green, D. M., & Swets, J. A. (1966). Signal detection theory and psychophysics. Wiley & Sons.

Hackley, S. A., Langner, R., Rolke, B., Erb, M., Grodd, W., & Ulrich, R. (2009). Separation of phasic arousal and expectancy effects in a speeded reaction time task via fMRI. Psychophysiology, 46(1), 163–171. 10.1111/j.1469-8986.2008.00722.x

Hackley, S. A., & Valle-Inclán, F. (1998). Automatic alerting does not speed late motoric processes in a reaction-time task. Nature, 391(6669), 786–788. 10.1038/35849

Hershenson, M. (1962). Reaction time as a measure of intersensory facilitation. Journal of Experimental Psychology, 63(3), 289–293. 10.1037/h0039516

James, T., Kula, B., Choi, S., Khan, S. S., Bekar, L. K., & Smith, N. A. (2021). Locus coeruleus in memory formation and Alzheimer’s disease. The European Journal of Neuroscience, 54(8), 6948–6959. 10.1111/ejn.15045

Jepma, M., Wagenmakers, E.-J., Band, G. P. H., & Nieuwenhuis, S. (2009). The effects of accessory stimuli on information processing: Evidence from electrophysiology and a diffusion model analysis. Journal of Cognitive Neuroscience, 21(5), 847– 864. 10.1162/jocn.2009.21063

Jiménez, E. C., Sierra-Marcos, A., Romeo, A., Hashemi, A., Leonovych, O., Bustos Valenzuela, P., Solé Puig, M., & Supèr, H. (2021). Altered Vergence Eye Movements and Pupil Response of Patients with Alzheimer’s Disease and Mild Cognitive Impairment During an Oddball Task. Journal of Alzheimer’s Disease, 82(1), 421–433. 10.3233/JAD-201301

Jordan, R. (2023). The locus coeruleus as a global model failure system. Trends in Neurosciences, 0(0). 10.1016/j.tins.2023.11.006

Joshi, S., & Gold, J. I. (2020). Pupil Size as a Window on Neural Substrates of Cognition. Trends in Cognitive Sciences, 24(6), 466–480. 10.1016/j.tics.2020.03.005

Joshi, S., & Gold, J. I. (2022). Context-dependent relationships between locus coeruleus firing patterns and coordinated neural activity in the anterior cingulate cortex. ELife, 11. 10.7554/eLife.63490

Joshi, S., Li, Y., Kalwani, R. M., & Gold, J. I. (2016). Relationships between Pupil Diameter and Neuronal Activity in the Locus Coeruleus, Colliculi, and Cingulate Cortex. Neuron, 89(1), 221–234. 10.1016/j.neuron.2015.11.028

Kloosterman, N. A., Meindertsma, T., van Loon, A. M., Lamme, V. A. F., Bonneh, Y. S., & Donner, T. H. (2015). Pupil size tracks perceptual content and surprise. The European Journal of Neuroscience, 41(8), 1068–1078. 10.1111/ejn.12859

Knapen, T., de Gee, J. W., Brascamp, J., Nuiten, S., Hoppenbrouwers, S., & Theeuwes, J. (2016). Cognitive and Ocular Factors Jointly Determine Pupil Responses under Equiluminance. PloS One, 11(5), e0155574. 10.1371/journal.pone.0155574

Krishnamurthy, K., Nassar, M. R., Sarode, S., & Gold, J. I. (2017). Arousal-related adjustments of perceptual biases optimize perception in dynamic environments. Nature Human Behaviour, 1(6), 1–11. 10.1038/s41562-017-0107

Lavín, C., San Martín, R., & Rosales Jubal, E. (2013). Pupil dilation signals uncertainty and surprise in a learning gambling task. Frontiers in Behavioral Neuroscience, 7, 218. 10.3389/fnbeh.2013.00218

Lewandowska, K., Gągol, A., Sikora-Wachowicz, B., Marek, T., & Fąfrowicz, M. (2019). Saying “yes” when you want to say “no” - pupil dilation reflects evidence accumulation in a visual working memory recognition task. International Journal of Psychophysiology, 139, 18–32. 10.1016/j.ijpsycho.2019.03.001

Lloyd, B., de Voogd, L. D., Mäki-marttunen, V., & Nieuwenhuis, S. (2023). Pupil size reflects activation of subcortical ascending arousal system nuclei during rest. ELife, 12, e84822. 10.7554/eLife.84822

Macmillan, N. A., & Creelman, C. D. (2005). Detection theory: A user’s guide (2^nd^ ed.). Lawrence Erlbaum Associates.

Murphy, P. R., Krkovic, K., Monov, G., Kudlek, N., Lincoln, T., & Donner, T. H. (2024). Individual Differences in Belief Updating and Phasic Arousal Are Related to Psychosis Proneness. BioRxiv, 2024-01. 10.1101/2024.01.14.575567

Murphy, P. R., O’Connell, R. G., O’Sullivan, M., Robertson, I. H., & Balsters, J. H. (2014). Pupil diameter covaries with BOLD activity in human locus coeruleus. Human Brain Mapping, 35(8), 4140–4154. 10.1002/hbm.22466

Murphy, P. R., Wilming, N., Hernandez-Bocanegra, D. C., Prat-Ortega, G., & Donner, T. H. (2021). Adaptive circuit dynamics across human cortex during evidence accumulation in changing environments. Nature Neuroscience, 24(7), 987–997. 10.1038/s41593-021-00839-z

Nassar, M. R., Rumsey, K. M., Wilson, R. C., Parikh, K., Heasly, B., & Gold, J. I. (2012). Rational regulation of learning dynamics by pupil-linked arousal systems. Nature Neuroscience, 15(7), 1040–1046. 10.1038/nn.3130

Okazawa, G., Sha, L., Purcell, B. A., & Kiani, R. (2018). Psychophysical reverse correlation reflects both sensory and decision-making processes. Nature Communications, 9(1), 3479. 10.1038/s41467-018-05797-y

Petersen, A., Petersen, A. H., Bundesen, C., Vangkilde, S., & Habekost, T. (2017). The effect of phasic auditory alerting on visual perception. Cognition, 165, 73–81. 10.1016/j.cognition.2017.04.004

Poe, G. R., Foote, S., Eschenko, O., Johansen, J. P., Bouret, S., Aston-Jones, G., Harley, C. W., Manahan-Vaughan, D., Weinshenker, D., Valentino, R., Berridge, C., Chandler, D. J., Waterhouse, B., & Sara, S. J. (2020). Locus coeruleus: A new look at the blue spot. Nature Reviews Neuroscience, 21(11), 644–659. 10.1038/s41583-020-0360-9

Posner, M. I., & Boies, S. J. (1971). Components of attention. Psychological Review, 78(5), 391–408. 10.1037/h0031333

Preuschoff, K., ’t Hart, B. M., & Einhäuser, W. (2011). Pupil Dilation Signals Surprise: Evidence for Noradrenaline’s Role in Decision Making. Frontiers in Neuroscience, 5, 115. 10.3389/fnins.2011.00115

Ranjbar-Slamloo, Y., & Fazlali, Z. (2019). Dopamine and Noradrenaline in the Brain; Overlapping or Dissociate Functions? Frontiers in Molecular Neuroscience, 12, 334. 10.3389/fnmol.2019.00334

Raz, A., & Buhle, J. (2006). Typologies of attentional networks. Nature Reviews Neuroscience, 7(5), 367–379. 10.1038/nrn1903

Ross, L. N., & Bassett, D. S. (2024). Causation in neuroscience: Keeping mechanism meaningful. Nature Reviews Neuroscience, 25(2), 81–90. 10.1038/s41583-023-00778-7

Sales, A. C., Friston, K. J., Jones, M. W., Pickering, A. E., & Moran, R. J. (2019). Locus Coeruleus tracking of prediction errors optimises cognitive flexibility: An Active Inference model. PLOS Computational Biology, 15(1), e1006267. 10.1371/journal.pcbi.1006267

Sara, S. J. (2009). The locus coeruleus and noradrenergic modulation of cognition. Nature Reviews Neuroscience, 10(3), 211–223. 10.1038/nrn2573

Schriver, B. J., Perkins, S. M., Sajda, P., & Wang, Q. (2020). Interplay between components of pupil-linked phasic arousal and its role in driving behavioral choice in Go/No-Go perceptual decision-making. Psychophysiology, 57(8), e13565. 10.1111/psyp.13565

Stahl, J., & Rammsayer, T. H. (2005). Accessory stimulation in the time course of visuomotor information processing: Stimulus intensity effects on reaction time and response force. Acta Psychologica, 120(1), 1–18. 10.1016/j.actpsy.2005.02.003

Tona, K.-D., Murphy, P. R., Brown, S. B. R. E., & Nieuwenhuis, S. (2016). The accessory stimulus effect is mediated by phasic arousal: A pupillometry study. Psychophysiology, 53(7), 1108–1113. 10.1111/psyp.12653

Tosserams, A., Bloem, B. R., Ehgoetz Martens, K. A., Helmich, R. C., Kessels, R. P. C., Shine, J. M., Taylor, N. L., Wainstein, G., Lewis, S. J. G., & Nonnekes, J. (2023). Modulating arousal to overcome gait impairments in Parkinson’s disease: How the noradrenergic system may act as a double-edged sword. Translational Neurodegeneration, 12(1), 15. 10.1186/s40035-023-00347-z

Totah, N. K., Neves, R. M., Panzeri, S., Logothetis, N. K., & Eschenko, O. (2018). The locus coeruleus is a complex and differentiated neuromodulatory system. Neuron, 99(5), 1055–1068. 10.1101/109710

Tsujimura, S., Wolffsohn, J. S., & Gilmartin, B. (2001). A linear chromatic mechanism drives the pupillary response. Proceedings. Biological Sciences, 268(1482), 2203– 2209. 10.1098/rspb.2001.1775

Urai, A. E., Braun, A., & Donner, T. H. (2017). Pupil-linked arousal is driven by decision uncertainty and alters serial choice bias. Nature Communications, 8, 14637. 10.1038/ncomms14637

van den Brink, R. L., Hagena, K., Wilming, N., Murphy, P. R., Calder-Travis, J., Finsterbusch, J., Büchel, C., & Donner, T. H. (2023). Brainstem arousal systems adaptively shape large-scale cortical interactions for flexible decision-making. BioRxiv, 2023-12. 10.1101/2023.12.05.570327

van den Brink, R. L., Murphy, P. R., & Nieuwenhuis, S. (2016). Pupil Diameter Tracks Lapses of Attention. PLOS ONE, 11(10), e0165274. 10.1371/journal.pone.0165274

Varazzani, C., San-Galli, A., Gilardeau, S., & Bouret, S. (2015). Noradrenaline and dopamine neurons in the reward/effort trade-off: A direct electrophysiological comparison in behaving monkeys. Journal of Neuroscience, 35(20), 7866–7877. 10.1523/JNEUROSCI.0454-15.2015

Wang, C.-A., & Munoz, D. P. (2015). A circuit for pupil orienting responses: Implications for cognitive modulation of pupil size. Current Opinion in Neurobiology, 33, 134–140. 10.1016/j.conb.2015.03.018

Waskom, M. L., Okazawa, G., & Kiani, R. (2019). Designing and Interpreting Psychophysical Investigations of Cognition. Neuron, 104(1), 100–112. 10.1016/j.neuron.2019.09.016

Wyart, V., Gardelle, V. de, Scholl, J., & Summerfield, C. (2012). Rhythmic fluctuations in evidence accumulation during decision making in the human brain. Neuron, 76(4), 847–858. 10.1016/j.neuron.2012.09.015

Yu, A. J., & Dayan, P. (2005). Uncertainty, neuromodulation, and attention. Neuron, 46(4), 681–692. 10.1016/j.neuron.2005.04.026

